# Silicone Wire Embolization-induced Acute Retinal Artery Ischemia and Reperfusion Model in Mouse: Gene Expression Provides Insight into Pathological Processes

**DOI:** 10.1101/2024.05.01.592074

**Authors:** Yuedan Wang, Ying Li, Jiaqing Feng, Chuansen Wang, Yuwei Wan, Bingyang Lv, Yinming Li, Hao Xie, Ting Chen, Faxi Wang, Ziyue Li, Anhuai Yang, Xuan Xiao

## Abstract

Acute retinal ischemia and ischemia-reperfusion injury are the primary causes of retinal neural cell death and vision loss in retinal artery occlusion (RAO). The absence of an accurate mouse model for simulating the retinal ischemic process has hindered progress in developing neuroprotective agents for RAO. We developed a unilateral pterygopalatine ophthalmic artery occlusion (UPOAO) mouse model using silicone wire embolization combined with carotid artery ligation. The survival of retinal ganglion cells and visual function were evaluated to determine the duration of ischemia. Immunofluorescence staining, optical coherence tomography, and haematoxylin and eosin staining were utilized to assess changes in major neural cell classes and retinal structure degeneration at two reperfusion durations. Transcriptomics was employed to investigate alterations in the pathological process of UPOAO following ischemia and reperfusion, highlighting transcriptomic differences between UPOAO and other retinal ischemia-reperfusion models. The UPOAO model successfully replicated the acute interruption of retinal blood supply observed in RAO. 60-minutes of Ischemia led to significant loss of major retinal neural cells and visual function impairment. Notable thinning of the inner retinal layer, especially the ganglion cell layer, was evident post-UPOAO. Temporal transcriptome analysis revealed various pathophysiological processes related to immune cell migration, oxidative stress, and immune inflammation during the non-reperfusion and reperfusion periods. A pronounced increase in microglia within the retina and peripheral leukocytes accessing the retina was observed during reperfusion periods. Comparison of differentially expressed genes (DEGs) between the UPOAO and high intraocular pressure models revealed specific enrichments in lipid and steroid metabolism-related genes in the UPOAO model. The UPOAO model emerges as a novel tool for screening pathogenic genes and promoting further therapeutic research in RAO.

## 1 Introduction

Retinal artery occlusion (RAO) is a severe ophthalmic disease characterized by a sudden interruption of blood flow in the retinal artery, leading to retinal ischemia [1]. Over 60% of RAO patients suffer from impaired vision, ranging from finger counting to complete vision loss [2]. Additionally, RAO patients face increased risk of cardiovascular and cerebrovascular events [3–5]. Retinal ischemia and hypoxia can lead to irreversible damage to retinal cells within 90 minutes [6]. This damage is primarily due to the apoptosis of retinal ganglion cells (RGCs), which is driven by inflammation and oxidative stress. Unfortunately, conservative treatments for RAO, such as ocular massage and hyperbaric oxygen therapy, provide limited therapeutic benefits [7]. Although thrombolysis is effective, its application is restricted by a narrow therapeutic window (typically within 4.5 hours) [8]. Furthermore, the restoration of blood flow to the ischemic area following thrombolysis can cause a sudden increase in tissue oxidative levels, resulting in ischemia-reperfusion injury (IRI) in the retina [9]. IRI is a common pathological condition that can induce inflammation, retinal tissue damage and visual impairment [10]. Therefore, it is crucial to develop an animal model that accurately simulates the pathological processes of RAO to thoroughly investigate the pathophysiological changes and explore potential neuroprotective treatments following retinal ischemia and reperfusion.

Various animal models have been employed to investigate the effects of retinal injury resulting from ischemia and subsequent reperfusion in RAO [6, 11, 12]. Based on the methods used for modelling, retinal ischemia-reperfusion models can be categorized into two groups: 1) intravascular occlusion models, including photochemical-induced thrombosis model and vascular intervention model; and 2) extravascular occlusion models, including central retinal artery ligation (CRAL) model, unilateral common carotid artery occlusion (UCCAO) model, and high intraocular pressure (HIOP) model. The photochemical-induced thrombosis model and vascular intervention model have been reported to be useful tools for assessing retinal injury after ischemia and reperfusion. However, both models are limited by the need for trained interventional radiologists and the need for advanced techniques [13, 14]. The CRAL model, which directly clips or ligates the central retinal artery, also presents drawbacks, such as potential optic nerve damage and difficulty in application to small animals such as mice, limiting its use in experiments with larger sample sizes [6]. The UCCAO model, which induces retinal hypoperfusion by occluding the unilateral common carotid artery (CCA), is more suitable for simulating chronic ischemic retinal disease than acute retinal ischemia [15, 16]. The HIOP model is widely utilized in mice for studying ischemia-reperfusion in acute primary angle-closure glaucoma (APACG) [17]. However, the mechanical compression of the retina caused by saline injection in this model may also lead to retinal damage, potentially influencing IRI research. Therefore, it is critically important to develop a simple and low-skill-required mouse model that can simulate acute retinal ischemia and reperfusion injury in RAO patients.

To better simulate the retinal ischemic process and possible IRI following RAO, we developed a novel vascular-associated mouse model called the unilateral pterygopalatine ophthalmic artery occlusion (UPOAO) model. In this model, we employed silicone wire embolization and carotid artery ligation to completely block the blood supply to the retina. We characterized the major classes of retinal neural cells and evaluated visual function following different durations of ischemia (30 minutes and 60 minutes) and reperfusion (3-days and 7-days) after UPOAO. Additionally, we utilized transcriptomics to investigate the transcriptional changes and elucidate the pathophysiological process of the UPOAO model after ischemia and reperfusion. Finally, we highlighted the transcriptomic differences between this model and other retinal ischemia-reperfusion models, including HIOP and UCCAO, revealing the unique pathological processes that closely resemble retinal IRI in RAO. The UPOAO model offers new insights into the mechanisms and pathways involved in ischemia and reperfusion studies of RAO, providing a foundation for studying protective strategies for ocular ischemic diseases.

## 2 Methods and materials

### 2.1 Animals

Eight-week-old male C57BL/6 mice were used in the experiments. Only male mice were used to exclude the potential influence of oestrogenic hormones. The mice were provided sufficient food and water and were maintained on a 12-hour dark-light cycle in a room with regulated temperature conditions. All the experimental procedures were designed and conducted according to the ethical guidelines outlined in the Association for Research in Vision and Ophthalmology (ARVO) Statement for the Use of Animals in Ophthalmic and Vision Research. The study protocol and methods received approval from the Experimental Animal Ethics Committee of Renmin Hospital of Wuhan University (approval number: WDRM-20220305A).

### 2.2 Preparation and surgical procedure of the UPOAO model

Eight-week-old male C57BL/6 mice weighing 20-25 grams were used in this study. An isoflurane-based anaesthesia system was used to induce and maintain general anaesthesia in the mice during surgical procedures. The mice were anaesthetized with a 1.5-2% concentration of isoflurane delivered in a mixture of nitrous oxide and oxygen via rubber tubing. Body temperature was maintained at 37 ± 0.5°C throughout the surgery. Surgical instruments were sterilized with 75% ethanol before use to ensure sterility.

We drew inspiration from the widely employed middle cerebral artery occlusion (MCAO) model [18, 19], commonly used in cerebral ischemic injury research, which guided the development of the UPOAO model. The mouse was gently positioned in a supine posture on a heating blanket, and its neck was exposed. The preparation involved depilation of the neck area, followed by skin disinfection, and finally, a midline incision along the neck was made. After separating the neck gland using two tweezers, careful blunt dissection was employed to separate the left CCA, internal carotid artery (ICA) and external carotid artery (ECA), avoiding the compression of nearby nerves and veins (**Fig. 1A**) (**Video 1**).

**Fig. 1.**
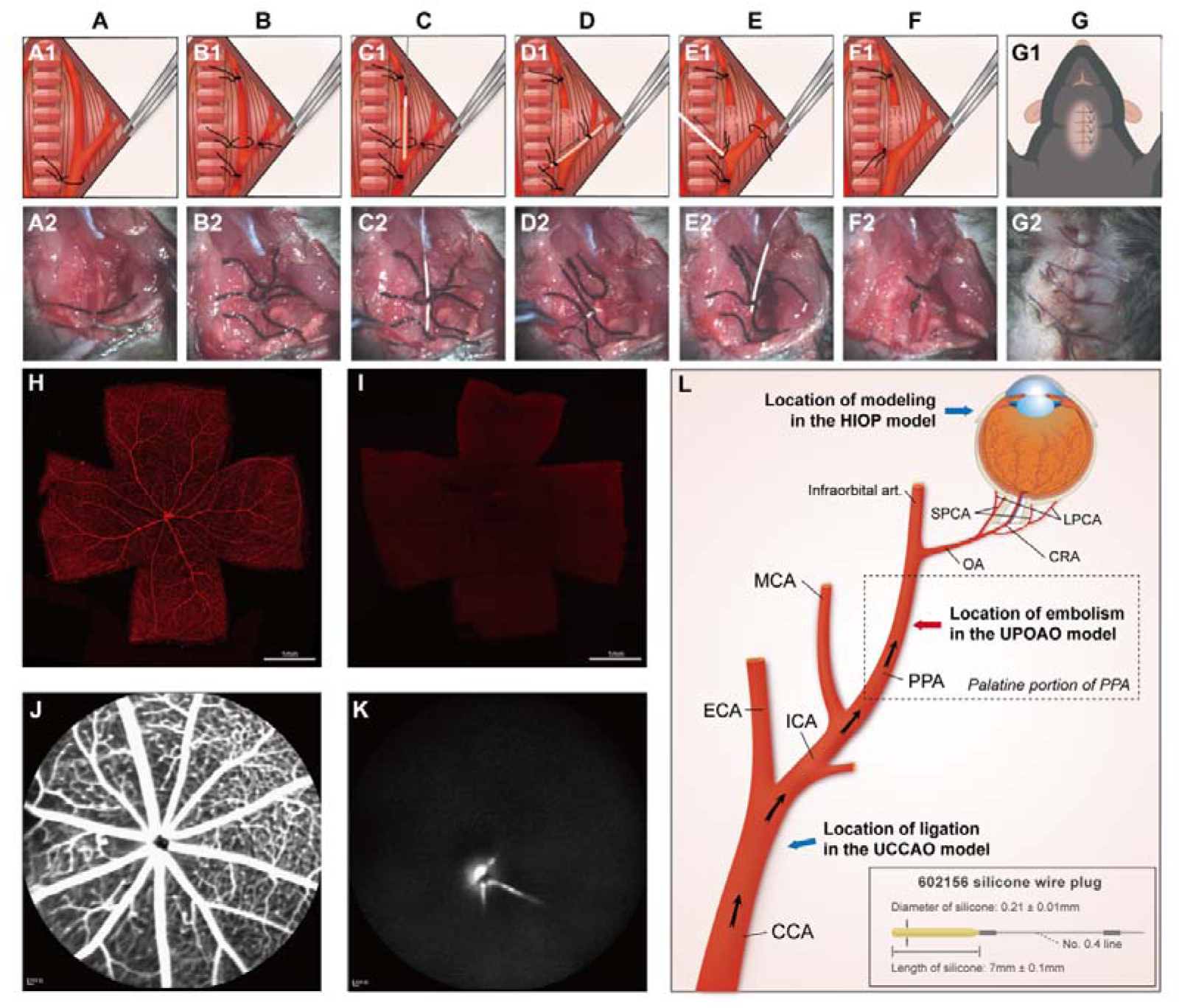
Modeling Procedure, Validation, and Cervical Artery Anatomy. (A1-G1) Schematic illustration of unilateral pterygopalatine ophthalmic artery occlusion (UPOAO). (A2-G2) Practical operation of UPOAO. (A1, A2) Blunt separation and exposure of the left cervical arteries. (B1, B2) Arterial suture ligation. (C1, C2) Insertion of the silicone wire embolus. The artery was incised to create a hole, and the silicone wire embolus was inserted. (D1, D2) Artery disconnection and movement of the silicone wire embolus. The artery was cut along the incision, and the silicone wire embolus was retracted and reinserted. (E1, E2) Removal of the silicone wire embolus and reperfusion. (F1, F2) Suture removal. The sutures at both ends of the disconnected vessel were knotted, and the other two sutures were removed. (G1, G2) Anatomic reduction and suturing of the skin. (H, I) Canavalin A label vasculature of the UPOAO mouse retina. The silicone wire embolus was inserted into artery before perfusing rhodamine-labeled canavalin A into the heart of UPOAO mouse. The sham eye served as an unpracticed control eye, while the UPOAO lateral eye represented the experimental eye. Retinal vessels in the sham eye (H) exhibited fluorescence filling, while retinal vessels in the UPOAO lateral eye (I) remained unfilled. Scale bar = 1 mm. (J and K) Fluorescein Fundus Angiography (FFA) performed before removing the silicone wire embolus from the UPOAO mouse. The vessels in the sham lateral retina (J) were perfused, while the lateral retinal perfusion in UPOAO (K) was delayed. (L) Schematic illustration of cervical artery anatomy and ocular blood supply. Embolization of the PPA resulted in ocular ischemia. The red arrow indicates the site of the silicone wire embolus occlusion. The silicone wire embolus used a type 602156 wire, extended to 7mm with a diameter of 0.21mm. The blue arrows indicate the modelling locations of the HIOP model and the UCCAO model, respectively. CCA: common carotid artery; ICA: internal carotid artery; ECA: external carotid artery; PPA: pterygopalatine artery; MCA: middle cerebral artery; Infraorbital art.: infraorbital artery; OA: ophthalmic artery; SPCA: short posterior ciliary artery; LPCA: long posterior ciliary artery; CRA: central retinal artery.

Then, the CCA and ICA were ligated with a 6-0 suture. A knot was secured using a 6-0 suture at the distal end of the ECA, and a slipknot was created at the proximal end (**Fig. 1B**). To preserve the reperfusion process and avoid bleeding and mortality during surgery, a silicone wire embolus was inserted through the ECA instead of through the CCA. Ophthalmic scissors were used to make a small inverted “V”-shaped incision between the two suture knots on the ECA. A specialized silicone wire embolus, measuring 7 ± 0.1 mm in length and 0.21 ± 0.1 mm in diameter, was inserted through the incision, directing it into the ECA and further into the CCA (**Fig. 1C**). Next, the ligation on the ICA was removed, and the ECA was cut at the point where the artery had been previously incised with scissors. Subsequently, the silicone wire embolus was retracted to the bifurcation of the CCA, rotated counterclockwise, and inserted into the ICA and further into the pterygopalatine artery (PPA) (**Fig. 1D1**). The silicone tail of the silicone wire embolus was positioned near the bifurcation of the CCA (**Fig. 1D2**), effectively obstructing the ophthalmic artery (OA) (**Video 2**). Then, the slipknot was fastened, and the skin was sutured. The mouse was free to move around during arterial embolization.

Following a predetermined ischemia embolization time, the silicone wire embolus was carefully removed from the PPA without massive haemorrhage (**Fig. 1E**). The CCA suture was removed, restoring arterial reperfusion (**Fig. 1F**) (**Video 3**). Then, the skin wound was sutured, and the mice were routinely fed during reperfusion (**Fig. 1G**).

### 2.3 Staining of retinal vessels

To clearly visualize the retinal vessels, rhodamine-labelled canavalin A was used for mouse heart perfusion. Each mouse was anaesthetized with 1% sodium pentobarbital and placed in the supine position on a foam board resting on a tray. The sternum was opened to expose the heart. A needle attached to the perfusion tube was inserted into the left ventricle via the apex of the mouse heart and secured. The right atrial appendage was dissected with scissors to allow blood outflow. Saline solution was perfused through the left ventricle using a pump system to drain blood from the arteries. Subsequently, the prepared solution of rhodamine-labelled canavalin A was injected into the mice and circulated throughout the mouse vessels for 30 minutes. The mice were then sacrificed, and their eyes were removed and immersed in paraformaldehyde (PFA) for one hour in the dark. A flat-mounted retina was obtained, and retinal images were captured using a Leica SP8 confocal microscope equipped with a 10× objective.

### 2.4 Quantification of RGCs and microglia

The mice were euthanized by cervical dislocation, and their eyes were fixed in 4% PFA for 60 minutes at room temperature (RT). Subsequently, the cornea and lens were excised, and the intact retina was isolated for retinal flat mounting. The retina was immersed in PBS supplemented with 5% bovine serum albumin (BSA) and 0.5% Triton X-100 for overnight blocking and then incubated with Brn3a antibody (Synaptic Systems, Germany) or Iba1 antibody (Wako, Japan). Following a two-day incubation with the primary antibody, the retina was gently washed three times with PBS. Subsequently, the retina was incubated with Alexa Fluor 594 (AntGene, Wuhan, China) in a light-protected cassette for two days. After three rinses, the retina was uniformly sectioned into a four-leaf clover morphology and flattened using a coverslip. Retinal flat mounts labelled with Brn3a were photographed to count using a fluorescence microscope (BX51, Olympus, Japan) and representative pictures were imaged using a Leica SP8 confocal microscope (Leica TCS SP8, Germany). Retinas labelled with Iba1 were imaged utilizing a fluorescence microscope (BX63; Olympus, Tokyo, Japan). Each quadrant of the retina was systematically subdivided into central, middle, and peripheral fields (distance from the optic nerve head: central field: 0.1 mm to 0.5 mm, middle field: 0.9 mm to 1.3 mm, and peripheral field: 1.7 mm to 2.1 mm). The surviving RGCs and activated microglia of 12 fields in each retina were quantified and averaged using ImageJ software (National Institutes of Health, USA).

### 2.5 Electroretinogram (ERG)

Mice were dark-adapted overnight before ERG, and all subsequent procedures were carried out in the dark. Before ERG, the mice were anaesthetized with 1% sodium pentobarbital via intraperitoneal injection, and their pupils were dilated. A subcutaneous electrode was inserted into the posterior cervical skin, a tail electrode was affixed to the posterior end of the mouse tail, and the corneal electrode was gently placed on the central corneal surface. A RetiMINER-C visual electrophysiological system (3VMED Co., Ltd., Shanghai, China) was used for recording electrical responses. The a-waves, b-waves, and oscillatory potentials (OPs) after flash stimuli of 0.01, 0.03, 0.1, 0.3, 1.0, 3.0, and 10.0 cd.s/m^2^ in scotopic adaptation were recorded. The amplitudes and implicit times of a-waves, b-waves, and OPs in response to various flash stimuli were analysed.

### 2.6 Optical Coherence Tomography Imaging (OCT) and Fluorescein Fundus Angiography (FFA)

The mice were placed on the platform of a Spectralis HRA + OCT device (Heidelberg Engineering, Heidelberg, Germany) for OCT imaging following anaesthesia. Pupil dilation was performed, and normal saline was applied regularly to maintain corneal moisture. The focal length of the device was adjusted until the mouse retina was clearly visible. The head of the mouse was gently repositioned to capture images of the peripheral fundus. The multiline mode was used to scan each layer of the retina, and four quadrants of view centred on the nipple in the upper left, lower left, upper right, and lower right were recorded. In this study, the total retinal thickness was manually segmented into three parts and encompassed the entire thickness from the nerve fibre layer to the photoreceptor layer. The ganglion cell complex (GCC) was defined as the combined thickness of the retinal nerve fibre layer (RNFL), ganglion cell layer (GCL), and inner plexiform layer (IPL). The inner nuclear layer (INL) and the outer plexiform layer (OPL) were combined for thickness analysis due to the difficulty in distinguishing between these layers. The remaining retinal layers, including the outer nuclear layer (ONL), inner segment/outer segment (IS/OS), and retinal pigment epithelium (RPE) layers, were measured together. The thickness of the retina at 1.5 papillary diameters (PD), 3.0 PD, and 4.5 PD from the centre of the optic disc was measured using the Heidelberg measuring tool.

For FFA, the mice were anaesthetized, and their pupils were dilated. Subsequently, fundus angiography was performed immediately following an intraperitoneal injection of sodium fluorescein. Images were acquired alternately for both eyes within 5 seconds of the injection.

### 2.7 Hematoxylin and eosin (HE)

Mouse eyes were fixed in FAS eyeball fixation solution (Service-bio, Wuhan, China) for 48 hours, followed by dehydration and subsequent embedding in paraffin. The paraffin blocks were trimmed parallel to the optic nerve to obtain 3-4 μm thick sections where the optic nerve was located. Six paraffin sections from each eyeball were stained with HE. Images were captured using an Olympus fluorescence microscope. The retinal thickness within a range of 200-1100 μm from the optic disc was measured using Image-Pro Plus 6.0 software.

### 2.8 Staining of Retinal Sections

For immunofluorescence, mouse eyes were fixed in 4% PFA solution and subjected to gradient dehydration using 10%, 20%, and 30% sucrose solutions. The following day, the eyes were embedded in optimum cutting temperature (OCT) compound (SAKURA, USA) and frozen at -80°C. Several 14-μm-thick frozen sections were obtained from each eyeball through the use of a freezing microtome (Leica, Wetzlar, Germany). The section surface was parallel to the optic nerve, and 3-6 frozen sections were mounted on a single slide. The slides were stored at -20°C until use.

All sections were blocked with 5% BSA and 0.5% Triton X-100 in PBS for 2 hours. Primary antibodies, as listed in **Table 1**, were diluted in PBS containing 5% BSA and 0.5% Triton X-100 and incubated at 4°C overnight. The following day, after three rinses, the sections were incubated with secondary antibodies at RT for 2 hours in a cassette, followed by three washes. The sections were then stained with DAPI (Service-bio, Wuhan, China) for 15 minutes. After the final three rinses, the frozen sections were sealed. A Leica SP8 instrument equipped with a 40× objective was used to photograph frozen sections near the optic nerve head. The fluorescence intensity of all slices of each eye was determined using ImageJ software.

**Table 1.**
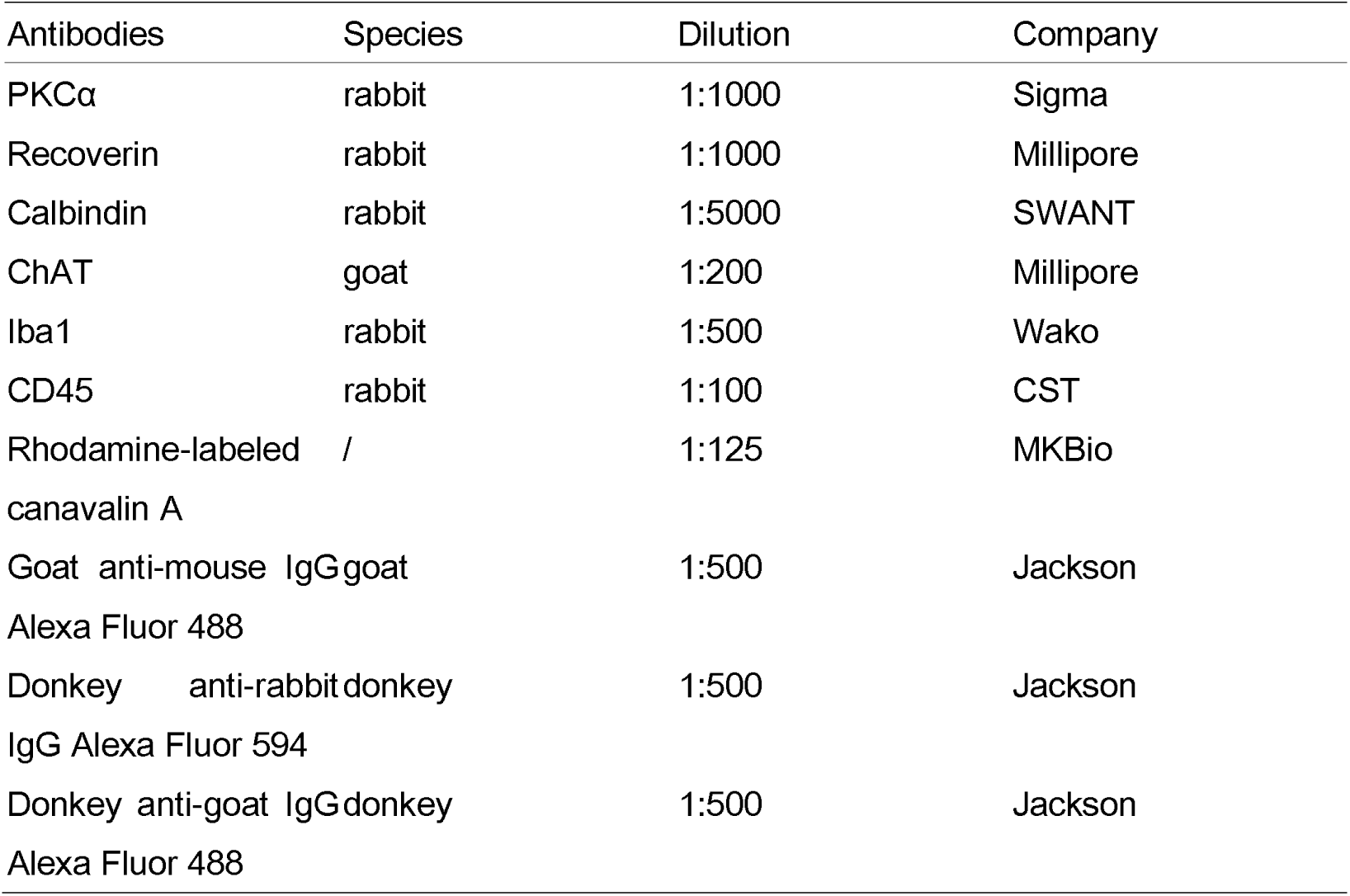
Antibodies Used in Staining of Flat-Mounted Retina and Sections.

### 2.9 Surgical procedure for the UCCAO model and the HIOP model

Animals for the UCCAO model were prepared and managed according to the procedure described for the UPOAO model. After blunt dissection, the left CCA was ligated with a 6-0 suture. After 60 minutes of ligation, the suture was removed, and the skin wound was subsequently closed with sutures. Mice were routinely fed during the reperfusion period, and the retinas of UCCAO mice were harvested 7-days post-surgery.

The HIOP model was generated based on previously established methods [20]. Briefly, mice were anaesthetized by intraperitoneal injection of 1% sodium pentobarbital. Pupils were dilated using tropicamide phenylephrine eye drops (0.5% tropicamide and 0.5% deoxyadrenaline hydrochloride, Santen Pharmaceutical) applied 5 minutes in advance. An insulin hypodermic needle attached to a silicone elastomer tube was inserted into the anterior chamber of the unilateral experimental eye. The IOP was elevated for 60 minutes by raising the height of the saline solution storage bag to 162 cm, after which the needle was removed to immediately restore the IOP to baseline levels. Iris whitening and loss of retinal red-light reflexes indicated retinal ischemia. After surgery, levofloxacin hydrochloride ophthalmic gel was applied to the eyes of the mice. Retinas from HIOP mice were collected 7-days post-surgery.

### 2.10 Transcriptome sequence and analysis

Retinal samples from UPOAO, HIOP, and UCCAO mice were extracted and promptly frozen in liquid nitrogen within 10 minutes of cervical dislocation. Total RNA extraction was performed according to the protocol outlined in the TRIzol Reagent manual (Life Technologies, CA, USA). RNA integrity and concentration were assessed using an Agilent 2,100 Bioanalyzer (Agilent Technologies, Inc., Santa Clara, CA, USA). The resulting RNA samples were then pooled and subjected to sequencing on the Illumina HiSeq3000 platform in a 150-bp paired-end read format. The raw RNA-sequencing (RNA-seq) reads were preprocessed and quantified using the featureCounts function in the SubReads package version 1.5.3, with default parameters.

RNA-seq analysis was conducted on samples from 5 mice in the non-perfusion group, 4 mice in the 3-days perfusion group, and 5 mice in the 7-days perfusion group of the UPOAO model. Additionally, retinas from both the 7-days HIOP experimental eyes and bilateral UCCAO eyes were collected (n=5) for analysis.

Data normalization and subsequent processing utilized the ‘limma’ package in R software (version 4.2.3) [21]. Volcano plots and Venn diagrams were generated for visualization to identify significantly differentially expressed genes (DEGs) and perform Gene Ontology (GO) annotation analysis. DEGs with a |log2-fold change (FC)| > 1 and adjusted p value < 0.05 were significantly differentially expressed. Enrichment analysis focused on DEGs with an adjusted p < 0.05 and an enriched gene count > 5. To visualize the expression patterns of significant DEGs, a heatmap was generated using the ‘pheatmap’ package. Gene Set Enrichment Analysis (GSEA) was conducted using the GSEA program (version 4.3.2) [22], employing the default gene set m-subset (mh.all.v2023.1.mm.symbols.gmt) to explore significant functional and pathway differences. Enriched pathways were classified based on criteria such as adjusted p value (< 0.05), false discovery rate (FDR) q value (< 0.25), and normalized enrichment score (|NES| > 1). Furthermore, protein ‒ protein interaction (PPI) analysis was performed using the Search Tool for the Retrieval of Interacting Genes (STRING) database (https://string-db.org/) [23]. Genes with a greater degree of protein-level interactions with others were further analysed using Cytoscape software (version 3.8.2) to generate a downstream PPI map. PPI pairings with an interaction score > 0.7 were extracted and visualized using Cytoscape 3.9.0 [24].

### 2.11 Real-time Quantitative polymerase chain reaction (RT-qPCR)

RNA was extracted from fresh mouse retinal samples and subjected to reverse transcription using HiScript lll RT SuperMix for qPCR (Vazyme, China) following the manufacturer’s instructions. RT‒qPCR analysis was conducted on a CFX instrument (Bio-Rad) using AceQ qPCR SYBR Green Master Mix (Vazyme, China). The PCR program comprised 40 cycles of 10 seconds at 95°C and 30 seconds at 60°C. Assays were performed in triplicate, and Ct values were normalized to beta-actin levels. Relative quantification of target gene expression was calculated using the 2-ΔΔCt method. The primer sequences are provided in **Table S1**.

### 2.12 Statistical Analysis

All the data were analysed using GraphPad Prism version 9.0 (GraphPad Software, San Diego, CA, USA). Each experimental group included a minimum of three biological replicates. T tests, including RGCs and microglia counting, OPs analysis, retinal thickness measurements via HE staining, and fluorescence intensity quantification, were used to compare the two groups. Two-way ANOVA (two-way analysis of variance) test was used for more than two different experimental groups, including ERG waves and retinal thickness measurements in OCT. All data are presented as the mean ± standard error of the mean (SEM), and the statistical graphs are shown as scatter bar graphs and line charts. P < 0.05 was considered to indicate statistical significance.

## 3 Results

### 3.1 Silicone wire embolus insertion interrupts retinal blood flow

To reproduce the retinal ischemic process and potential IRI in RAO, we established a mouse model of UPOAO by combining silicone wire embolization with carotid artery ligation. Model validation was conducted through a series of experiments, with detailed descriptions provided in the Materials and Methods section (**Fig. 1A-G**). To confirm blood flow disruption, we performed cardiac perfusion with fluorescently labelled lectin and FFA.

Rhodamine-labelled canavalin A was used during in vivo perfusion with silicone wire embolus, and both eyes were evaluated. The sham eye exhibited normal blood perfusion in the retina, while the experimental eye showed no perfusion (**Fig. 1H, I**). FFA revealed delayed and limited perfusion in the experimental lateral retina, primarily near the optic disc (**Fig. 1J, K**). These findings indicate that the insertion of the silicone wire embolus effectively impaired blood flow to the retina.

### 3.2 60-minute ischemia in UPOAO damage retinal structure and function

To investigate the optimal ischemic duration, the mice were subjected to ischemic periods of 30 and 60 minutes, followed by reperfusion periods of 3-days and 7-days. Retinal structure and visual function were assessed through flat-mounted retina analysis and flash ERG.

In the 30-minute ischemia group, no significant RGCs death was observed after 3-days (**Fig. 2A, B**) or 7-days of reperfusion (**Fig. 2C, D**). However, after 60 minutes of ischemia followed by either 3-days or 7-days of reperfusion, reductions in RGCs density were evident (**Fig. 2E-H**). ERG results showed no statistical difference in amplitudes between bilateral eyes after 3- and 7-days of 30-minutes reperfusion (**Fig. 3A-C**). However, the b-wave amplitude notably decreased after 60 minutes of ischemia and 3-days of reperfusion (**Fig. 3D**). By 7-days of reperfusion, the b-wave amplitude had halved compared to that of the sham eyes (**Fig. 3E**). The appearance times of the a-waves and b-waves in the experimental and control eyes remained consistent across all four groups (**Fig. S1**). Additionally, OPs have also been used to sensitively monitor retinal ischemic effects by detecting changes before any alterations in b-waves occur [25–27]. The amplitudes of OPs in the experimental eyes, particularly in the 60-minute ischemia and 7-days reperfusion group, significantly decreased to less than 50% of those in the sham eyes (**Fig. 3F, G**). Based on RGCs survival and changes in ERG waves, we determined that a 60-minute ischemic duration is optimal for the UPOAO model.

**Fig. 2.**
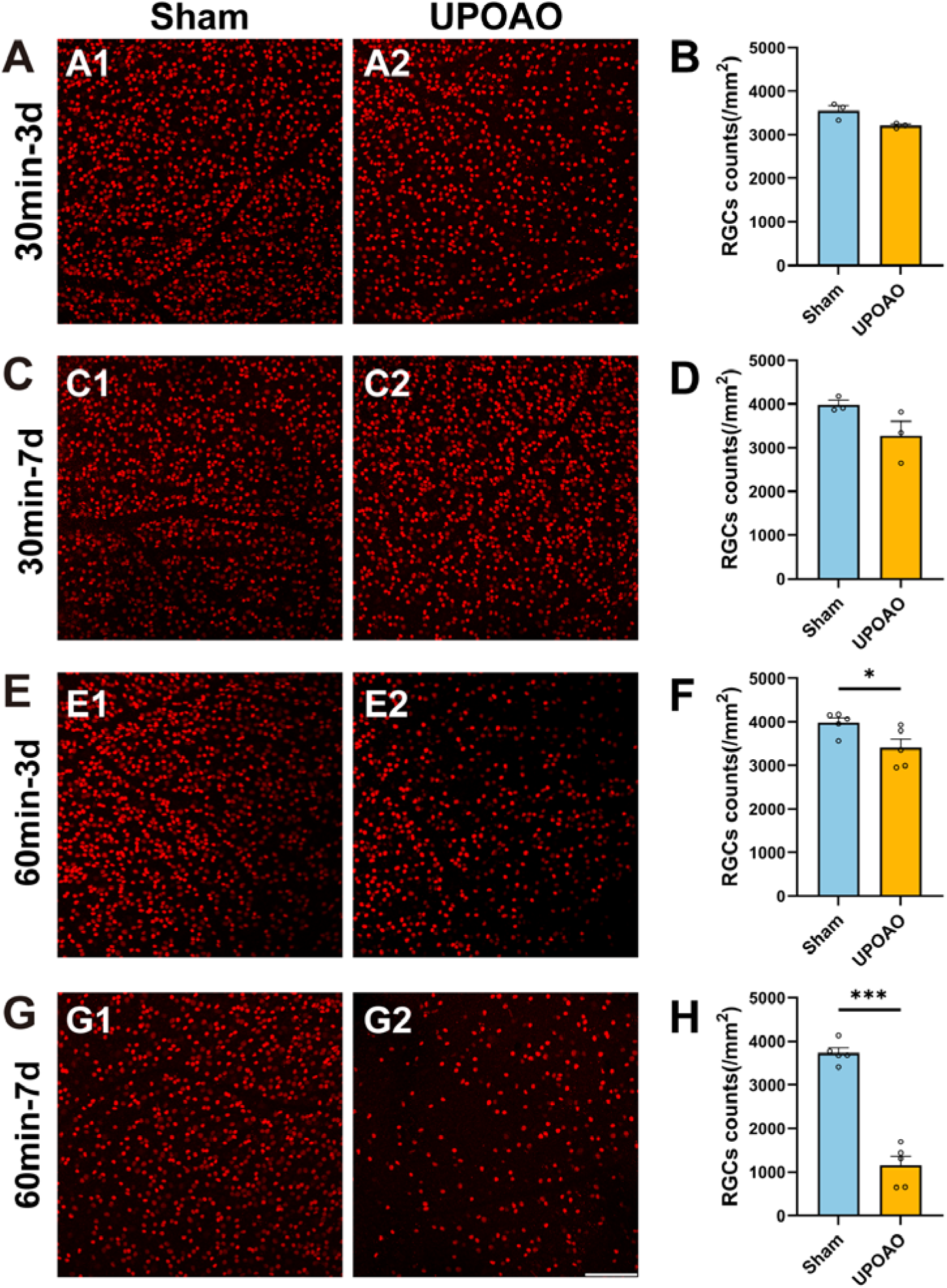
Staining and Quantification of Retinal Ganglion Cells (RGCs) at Different Ischemia and Reperfusion Times. Flat-mounted retina RGCs were labeled with Brn3a staining. (A, B) Representative pictures of the peripheral field (A), and quantification of surviving RGCs in all fields (B) in the 30-minute ischemia and 3-days reperfusion group. n = 3. (C, D) Representative pictures of the peripheral field (C), and quantification of surviving RGCs in all fields (D) in the 30-minute ischemia and 7-days reperfusion group. n = 3. (E, F) Representative pictures of the peripheral field (E), and quantification of surviving RGCs in all fields (F) in the 60-minute ischemia and 3-days reperfusion group. n = 5. (G, H) Representative pictures of the peripheral field (G), and quantification of surviving RGCs in all fields (H) in the 60-minute ischemia and 7-days reperfusion group. n = 5. The results showed a significant loss of RGCs after 60 minutes of ischemia. Data were presented as means ± s.e.m, *: p<0.05, **: p<0.01, ***: p<0.001, ****: p<0.0001, t-test. Scale bar = 100 μm.

**Fig. 3.**
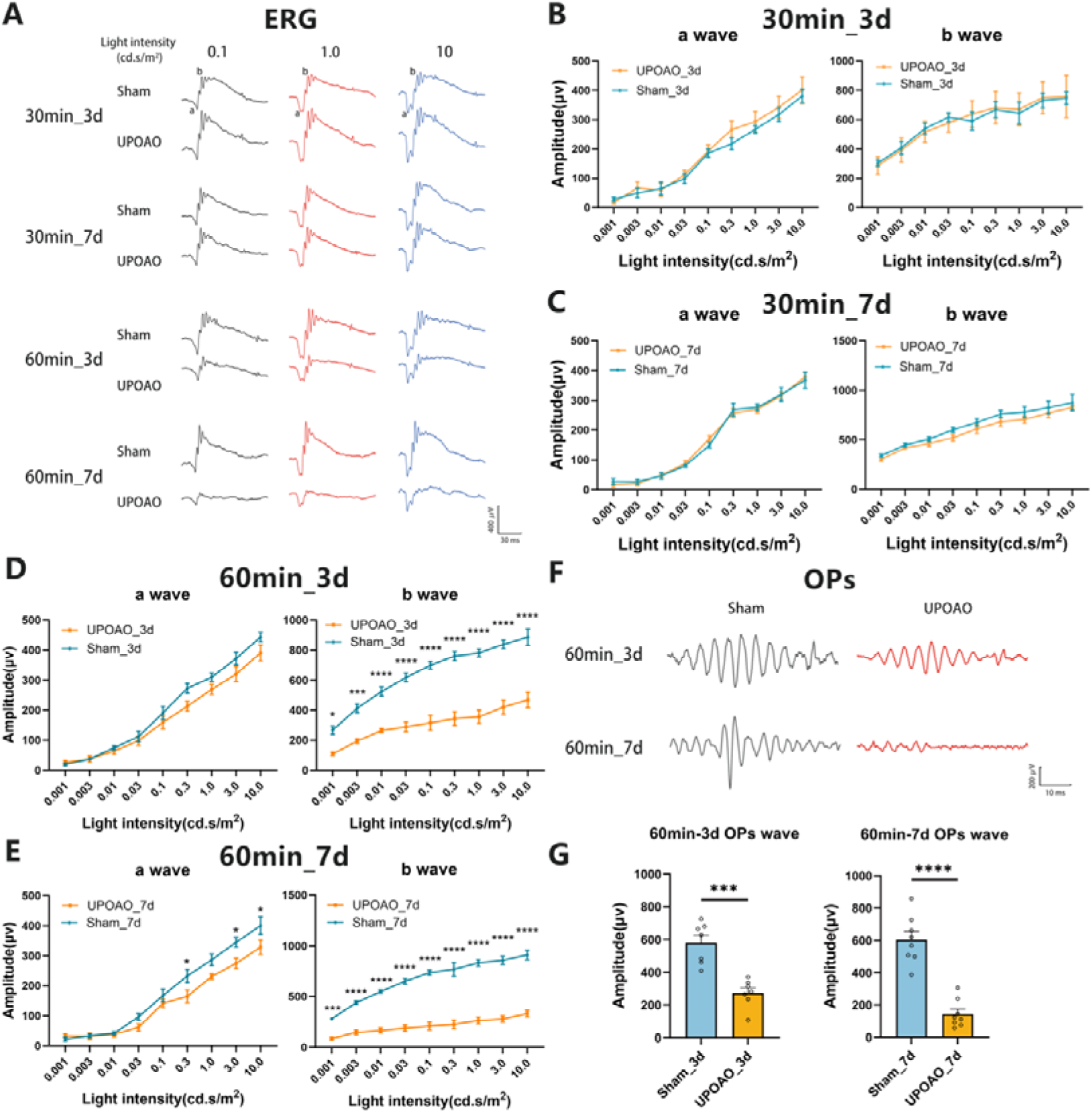
Comparison of Electroretinographic (ERG) Dark-Adapted Responses at Different Ischemia and Reperfusion Times. Following the evaluation of surviving RGCs, visual function in sham and UPOAO experimental eyes at various ischemia and reperfusion times was assessed using ERG. (A) Representative waveforms in the four groups at the stimulus light intensities of 0.1, 1.0, and 10.0 cd.s/m2, respectively. (B) Quantification of a-wave and b-wave amplitudes in the 30-minute ischemia and 3-days reperfusion group. n = 5. (C) Quantification of a-wave and b-wave amplitudes in the 30-minute ischemia and 7-days reperfusion group. n = 5. (D) Quantification of a-wave and b-wave amplitudes in the 60-minute ischemia and 3-days reperfusion group. n = 7. (E) Quantification of a-wave and b-wave amplitudes in the 60-minute ischemia and 7-days reperfusion group. n = 8. Dark-adapted responses showed almost similar a-wave amplitudes but significantly decreased b-wave amplitudes in the 60-minute ischemia groups. The amplitudes of b-waves declined at 3-days and even more prominently at 7-days. (F, G) Representative OPs and quantification of amplitudes in the 60-minute ischemic groups. n = 7 in the 3-days reperfusion group; n = 8 in the 7-days reperfusion group. The amplitudes of OPs decreased significantly at 7-days reperfusion. The decline in b-waves and OPs along with the loss of RGCs, supports the selection of a 60-minute ischemic duration as an appropriate choice. Data were presented as means ± s.e.m, *: p<0.05, **: p<0.01, ***: p<0.001, ****: p<0.0001, two-way ANOVA test for a-waves and b-waves; paired t-test for OPs.

### 3.3 Evaluation of retinal thickness in UPOAO through OCT and HE

To assess retinal thickness non-invasively in the UPOAO model, we conducted OCT imaging during 3-days and 7-days reperfusion periods. Following a 3-days reperfusion period, no significant changes were observed in the GCC, INL + OPL, and ONL + IS/OS + RPE layers at 1.5, 3, and 4.5 PD from the optic disc (**Fig. 4B, Fig. S2A, B**). However, the total retinal thickness decreased at 4.5 PD (**Fig. 4C**). After 7-days of reperfusion, the total retinal thickness decreased at 1.5 PD, 3 PD, and 4.5 PD, primarily due to inner retinal thinning (**Fig. 4D-F, Fig. S2C, D**).

**Fig. 4.**
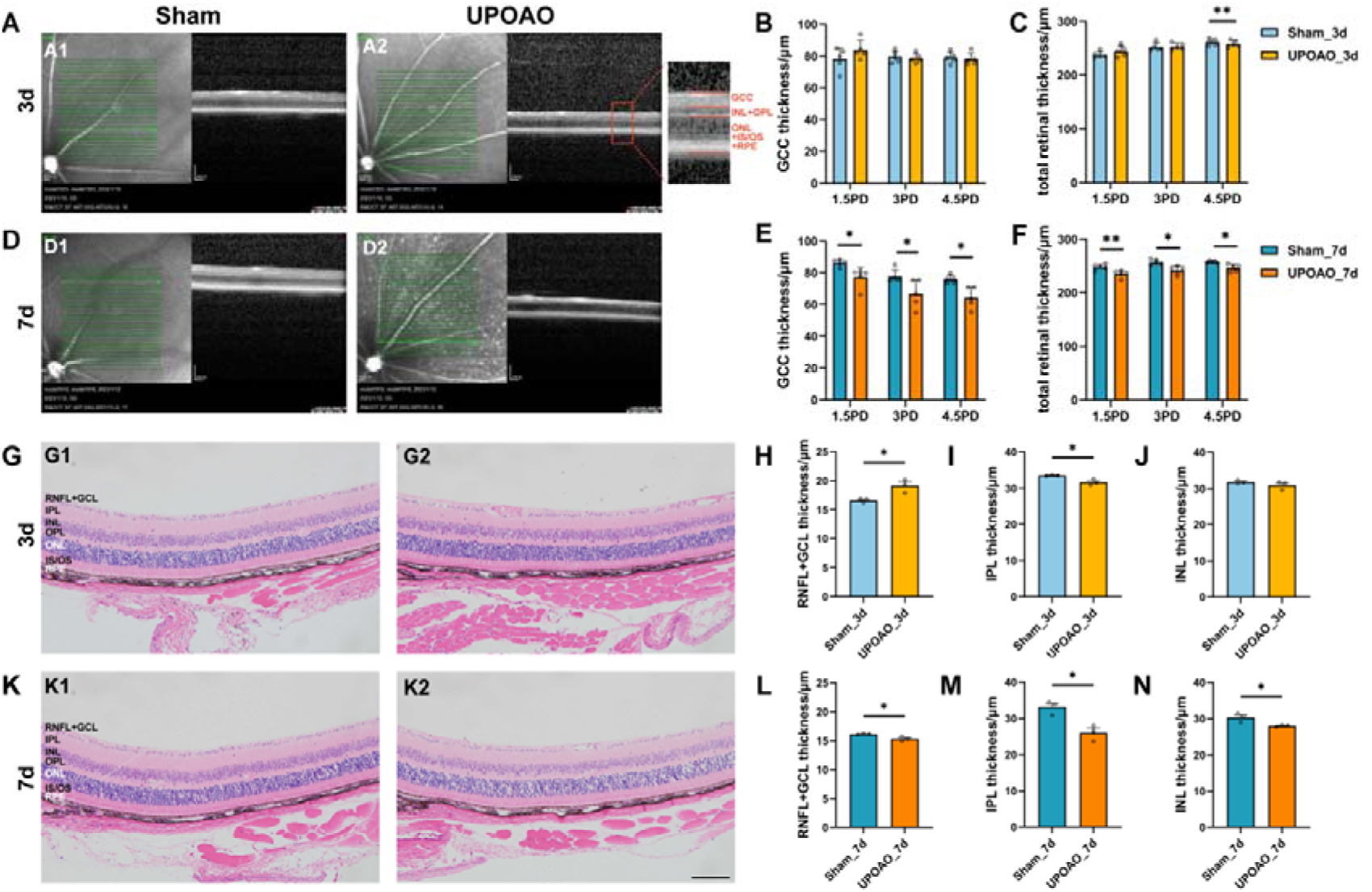
Changes in Retina Morphology in the UPOAO Model. (A) Representative OCT images of the mouse retina at 3-days. The Green lines indicate the OCT scan area starting from the optic disc. Local magnification and layering are annotated. GCC: ganglion cell complex, including RNFL, GCL and IPL layers. (B and C) Quantification of GCC and total retinal thickness at 3-days. The thickness of the GCC and the entire retina in OCT was measured and compared at distances of 1.5 PD, 3.0 PD and 4.5 PD from the optic disc, respectively. n = 5. (D) Representative OCT images of the mouse retina at 7-days. (E and F) Quantification of GCC and total retinal thickness at 7-days. n = 5. (G) Representative HE images of the mouse retina at 3-days. (H, I and J) Quantification of NFL + GCL, IPL, and INL thickness at 3-days. Retinal thickness in HE was measured near the optic nerve head and compared. n = 3. (K) Representative HE images of the mouse retina at 7-days. (L, M and N) Quantification of NFL + GCL, IPL, and INL thickness at 7-days. n = 3. Data were presented as means ± s.e.m, *: p<0.05, **: p<0.01, ***: p<0.001, ****: p<0.0001, two-way ANOVA test in OCT and paired t-test in HE. PD: papillary diameters; RNFL: retinal nerve fiber layer; GCL: ganglion cell layer; IPL: inner plexiform layer; INL: inner nuclear layer; OPL: outer plexiform layer; ONL: outer nuclear layer; IS: inner segment; OS outer segment; RPE: retinal pigment epithelium. Scale bar = 100 μm.

For a detailed analysis of inner retinal thickness, we extracted UPOAO mouse eyes and stained them with HE. The thickness of the RNFL + GCL increased at 3-days of reperfusion (**Fig. 4H**), followed by a decrease at 7-days of reperfusion (**Fig. 4L**). IPL thickness decreased at both 3-days and 7-days of reperfusion (**Fig. 4I, M**), while INL thickness decreased only at 7-days of reperfusion (**Fig. 4N**). We hypothesized that the initial swelling of the GCL in response to acute ischemia and hypoxia led to early thickening, followed by gradual thinning due to tissue dysfunction. This hypothesis was supported by the observation of RGCs loss and reduced GCC thickness evaluated by OCT at 7-days (**Fig. 4E**).

We observed by OCT that changes in retinal thickness became noticeable at 3-days post-UPOAO and exhibited a significant decrease at 7-days. HE results further revealed that alterations in inner retinal thickness constituted a substantial portion of the total retinal thickness during the early reperfusion stage after UPOAO.

### 3.4 Survival of retinal neural cells in UPOAO

Given that mouse retinal function heavily relies on rod cells and that scotopic (low light) vision is primarily governed by rod photoreceptors, the observed reductions in b-wave and OPs-wave amplitudes indicate impaired synaptic transmission within the inner retina [28]. Our OCT and HE findings suggested structural injuries within the inner retina. Notably, the thickness of the outer retinal layers in OCT remained unchanged. To further explore these observations, we conducted immunofluorescence staining to investigate alterations in major retinal cell types, primarily focusing on bipolar cells (BCs), photoreceptor cells, horizontal cells (HCs), and cholinergic amacrine cells.

#### 3.4.1 BCs loss and photoreceptor cells survival

PKCα serves as a marker delineating BCs, with their cell bodies primarily located in the outermost part of the INL, axonal terminals extending into the innermost part of the IPL, and dendrites confined to the OPL (**Fig. 5A, B**). In the experimental UPOAO eyes, BCs did not show significant changes at the 3-days reperfusion mark but exhibited dramatic alterations by the 7-days reperfusion period (**Fig. 5A2, B2**). In particular, the immunostaining density of BCs dendritic and axonal arbors notably decreased at the 7-days reperfusion mark. BCs account for approximately 40% of INL cells in mice, and their somata and axonal processes form a substantial part of the INL and IPL, consistent with the thinning of the INL observed in both OCT and HE at 7-days (**Fig. 4E, N**). Notably, the decrease in the number of PKCα^+^ BCs at the 7-days reperfusion point coincided with the decrease in the b-waves amplitude in ERG (**Fig. 3E**), indicating functional interplay between BCs and other cells.

**Fig. 5.**
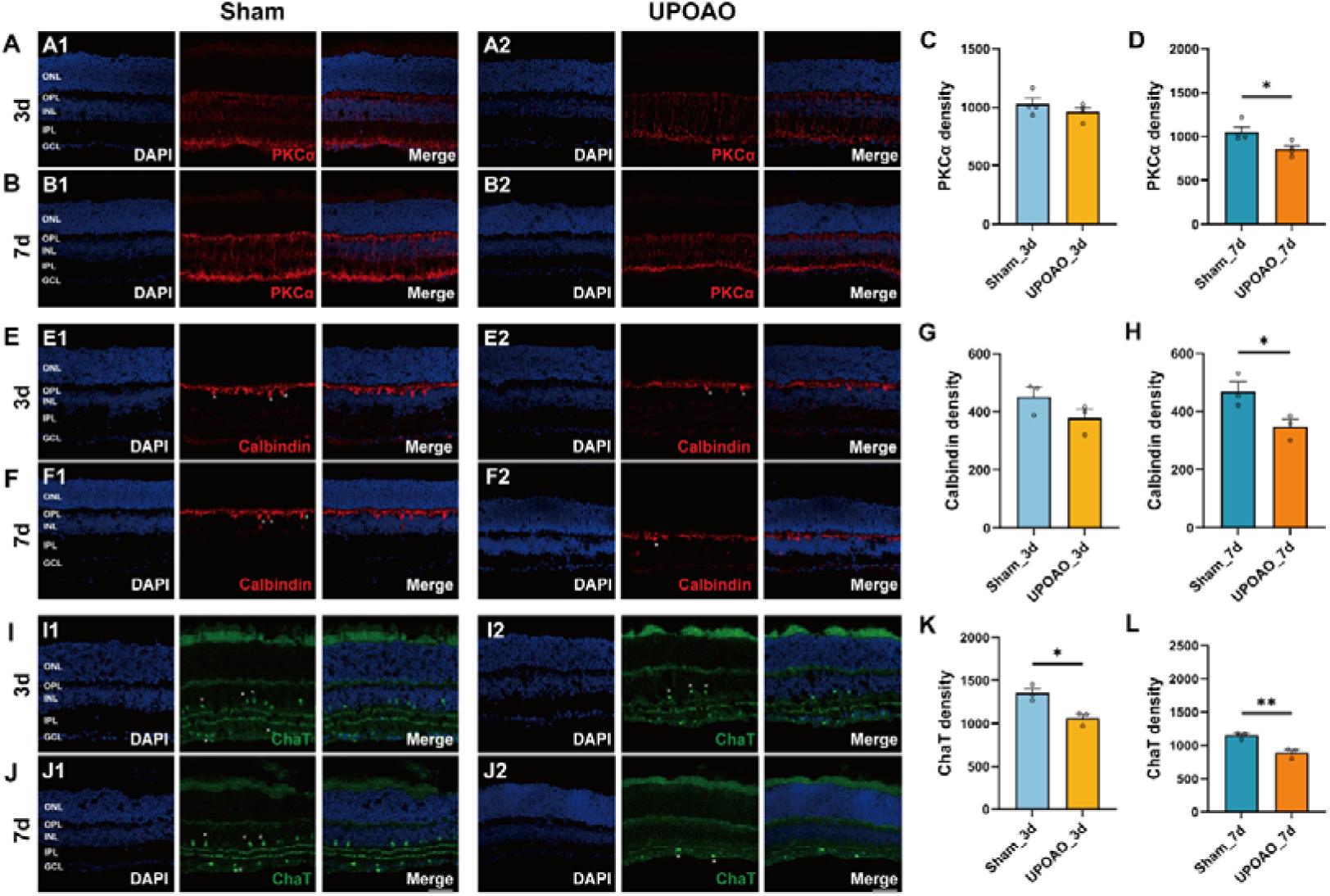
Changes in Bipolar Cells, Horizontal Cells, and Cholinergic Amacrine Cells in UPOAO Mice. (A) Representative images of mouse retina co-stained with DAPI and PKCα at 3-days (B) Representative images of mouse retina co-stained with DAPI and PKCα at 7-days. (C) Quantification of PKCα fluorescence density at 3-days. (D) Quantification of PKCα fluorescence density at 7-days. n = 4. (E) Representative images of mouse retina co-stained with DAPI and Calbindin at 3-days. (F) Representative images of mouse retina co-stained with DAPI and Calbindin at 7-days. Horizontal cell somata are indicated by white asterisks. (G) Quantification of Calbindin fluorescence density at 3-days. (H) Quantification of Calbindin fluorescence density at 7-days. n = 3. (I) Representative images of mouse retina co-stained with DAPI and ChAT at 3-days. (J) Representative images of mouse retina co-stained with DAPI and ChAT at 7-days. Cholinergic amacrine cell somata are indicated by white asterisks. (K) Quantification of ChAT fluorescence density at 3-days. (L) Quantification of ChAT fluorescence density at 7-days. n = 3. Data were presented as means ± s.e.m, *: p<0.05, **: p<0.01, ***: p<0.001, t-test. Scale bar = 50 μm.

Recoverin marks the somata and outer segments of photoreceptor cells and is localized within the ONL (**Fig. S3A, B**). Interestingly, recoverin-positive photoreceptors remained relatively stable throughout the reperfusion periods, which aligns with the unchanged thickness of the outer retinal layers observed in OCT (**Fig. S2C, D**).

#### 3.4.2 HCs and cholinergic amacrine cells loss

We evaluated HCs and cholinergic amacrine cells using calbindin and ChAT immunostaining, respectively. Calbindin immunostaining highlighted the somata of HCs within the INL, with terminal axon connections linearly positioned within the OPL (**Fig. 5E, F**). In the UPOAO experimental eyes, no significant change in the number of HCs was observed during the 3-days reperfusion period, while a notable reduction was observed after 7 days. By 7-days, a significant decrease in cell body numbers, a considerable reduction in axon density, and disrupted linear connections were observed (**Fig. 5F2**).

ChAT^+^ cell bodies were located in the GCL and INL, with dendrites forming two narrow stratified bands within the IPL. The immunofluorescence density of ChAT^+^ amacrine cells decreased notably after 3 days and even more prominently after 7 days (**Fig. 5I, J**). The fluorescence intensity of the two bands within the IPL markedly decreased and was nearly invisible after 7 days (**Fig. 5J2**).

### 3.5 Time course transcriptome analysis revealed features of different reperfusion periods in UPOAO

To explore the pathophysiological processes of UPOAO, we extracted retinas from both eyes for transcriptome sequencing. The samples included retinas from eyes subjected to 60 minutes of ischemia without reperfusion, 60 minutes of ischemia followed by 3-days of reperfusion, and 60 minutes of ischemia followed by 7-days of reperfusion, with the sham-treated eyes serving as controls.

In the non-perfusion group, 215 genes were upregulated and 204 genes were downregulated (**Fig. 6A**). GO enrichment analysis revealed that the DEGs were related to leukocyte migration, epidermis development, myeloid leukocyte migration and other cell migration pathways (**Fig. 6B**). The heatmap and box showed the upregulated and downregulated genes involved in immune cell migration-related pathways during the non-perfusion period in UPOAO (**Fig. 6C**). High-connectivity DEGs (‘hub genes’), such as Dusp1 and Fos, were also enriched in these pathways (**Fig. S4A**). GSEA also showed similar results (**Fig. S4B-D**). These results suggested that in the early stage of retinal ischemic injury, leukocytes from the microvasculature may infiltrate retinal tissue. More experimental validation will be performed to confirm this hypothesis.

**Fig. 6.**
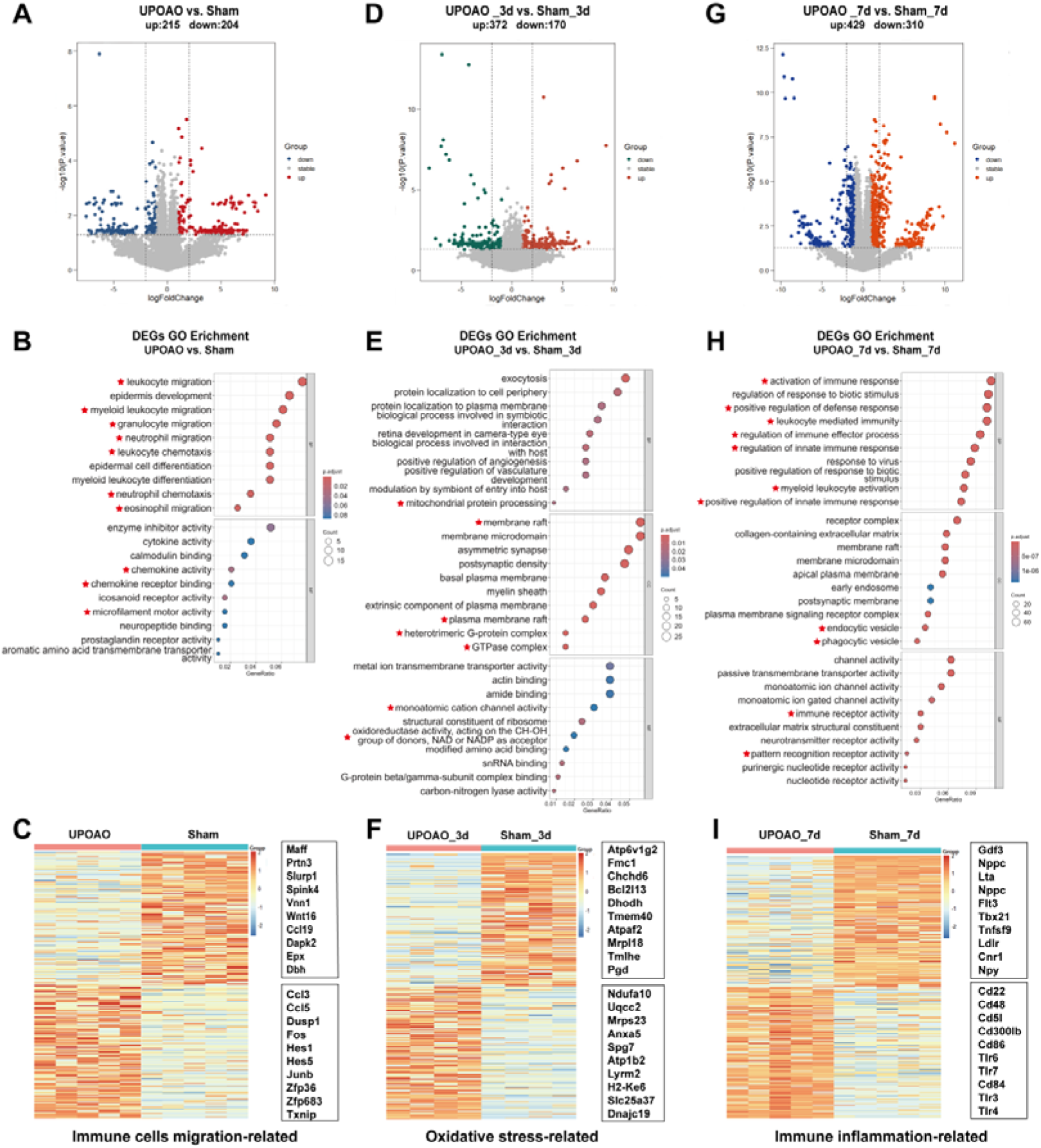
Transcriptomic Features at Different Times of Reperfusion. RNA-seq evaluation was performed at 0-day, 3-days, and 7-days reperfusion periods in UPOAO revealing enrichment in pathways related to immune cells migration, oxidative stress, and immune inflammation. (A, D, G) Volcano plots display differential expression genes (DEGs) between UPOAO and sham eyes in the no-perfusion group (A), 3-days perfusion group (D), and 7-days perfusion group (G), respectively. Red dots: significant upregulated genes, green and blue dots: significant downregulated genes, grey dots: stable expressed genes, adjusted P < 0.05. log2FC = 1. (B, E, H) Gene ontology (GO) analysis of differential genes in the non-perfusion group (B), 3-days group (E), and 7-days group (H). The DEGs between UPOAO and sham eyes were enriched in pathways associated with immune cells migration (0d), oxidative stress (3d), and immune inflammation (7d), respectively. (C, F, I) Heatmap displaying the top 100 downregulated and top 100 upregulated DEGs between UPOAO and sham at non-perfusion, 3-days, and 7-days reperfusion, respectively. The box represents the genes related to immune cells migration (C), oxidative stress (F), and immune inflammation (I). The ranking was determined by the magnitude of fold change. In each heatmap, the upper box represents the top 10 downregulated genes, while the lower box represents the top 10 upregulated genes.

In the 3-days reperfusion group, 372 genes were upregulated, and 170 genes were downregulated (**Fig. 6D**). GO enrichment analysis revealed that the DEGs at 3-days after reperfusion were related to energy metabolism, mitochondrial regulation, and oxidative stress pathways (**Fig. 6E**). The heatmap and box displayed the upregulated and downregulated genes involved in oxidative stress-related pathways at the 3-days reperfusion stage in UPOAO, including hub genes such as Mrpl18 and Mrps23 (**Fig. 6F, Fig. S5A**). GSEA revealed changes in pathways related to the binding membrane of organelles, the cell surface and the plasma membrane protein complex (**Fig. S5B**). Furthermore, 44 overlapping genes between the mitochondrial genes and DEGs at 3-days post-UPOAO were mainly related to mitochondrial transport and the mitochondrial tricarboxylic acid cycle (**Fig. S5C, D**). These results suggested that mitochondria and metabolism play significant roles in IRI.

In the 7-days reperfusion group, 429 genes were upregulated, and 310 genes were downregulated (**Fig 6G**). The DEGs at this stage were enriched in pathways related to immune effector processes, positive regulation of responses to external stimuli and inflammation (**Fig. 6H**). A heatmap was generated to show the upregulated and downregulated genes of immune inflammation-related pathways at 7-days after reperfusion in UPOAO, and most hub genes, such as Cd86, Cd48, Tlr4, and Tlr6 were also enriched in these pathways (**Fig. 6I, Fig. S6A**). GSEA also showed similar results (**Fig S6B**). Moreover, RT‒qPCR confirmed the upregulation of immune inflammation-related gene expression (**Fig. S7**). Significantly, 112 genes that overlapped between immune-related genes and DEGs at 7-days post-UPOAO were primarily associated with immune receptor activity and phagocytic vesicles (**Fig. S6C, D**). These results underscore the predominant role of immune responses during this stage.

To explore the associations between changes observed at 3-days and 7-days after reperfusion, we analyzed the DEGs and identified 17 overlapping genes (**Fig. S8A**), which were mainly associated with the nitric oxide biosynthetic process (**Fig. S8B**). Our findings suggest that the immune inflammatory response observed after 7-days of reperfusion may represent the cumulative effect of the acute oxidative stress response during the initial 3-days reperfusion period.

### 3.6 Leukocyte Infiltration and Microglial Activation

Time course transcriptome analysis underscored the predominant role of immune inflammatory responses during the reperfusion stage. Previous reports have indicated that the inflammatory response in retinal ischemia-reperfusion injury (RIRI) is orchestrated by peripheral immune cells and resident immune cells within the retina [29, 30]. In our study, we demonstrated the presence of leukocyte infiltration and microglial activation in the UPOAO model. Immunofluorescence staining for CD45 was performed on retinas at different time points post modelling to visualize the distribution and quantity of white blood cells (**Fig. 7A-D**). Laser confocal Z-plane projections confirmed minimal leukocyte infiltration in the retinas of sham eyes, which showed that CD45^+^ cells in the vascular lumen likely represented patrolling cells (**Fig. 7A**). A significant increase in CD45^+^ cells was observed in UPOAO model retinas after 1 day (**Fig. 7E**), with a progressive increase in quantity over time (**Fig. 7F, G**). Notably, the majority of CD45^+^ cells in UPOAO model retinas at 1-day displayed a morphology similar to that of vascular leukocytes (**Fig. 7B**), suggesting a large influx of peripheral white blood cells into the retinal tissue. In contrast, CD45^+^ cells at 3-days and 7-days of reperfusion exhibited a combination of amoeboid and branched morphologies (**Fig. 7C, D**), indicative of activated states.

**Fig. 7.**
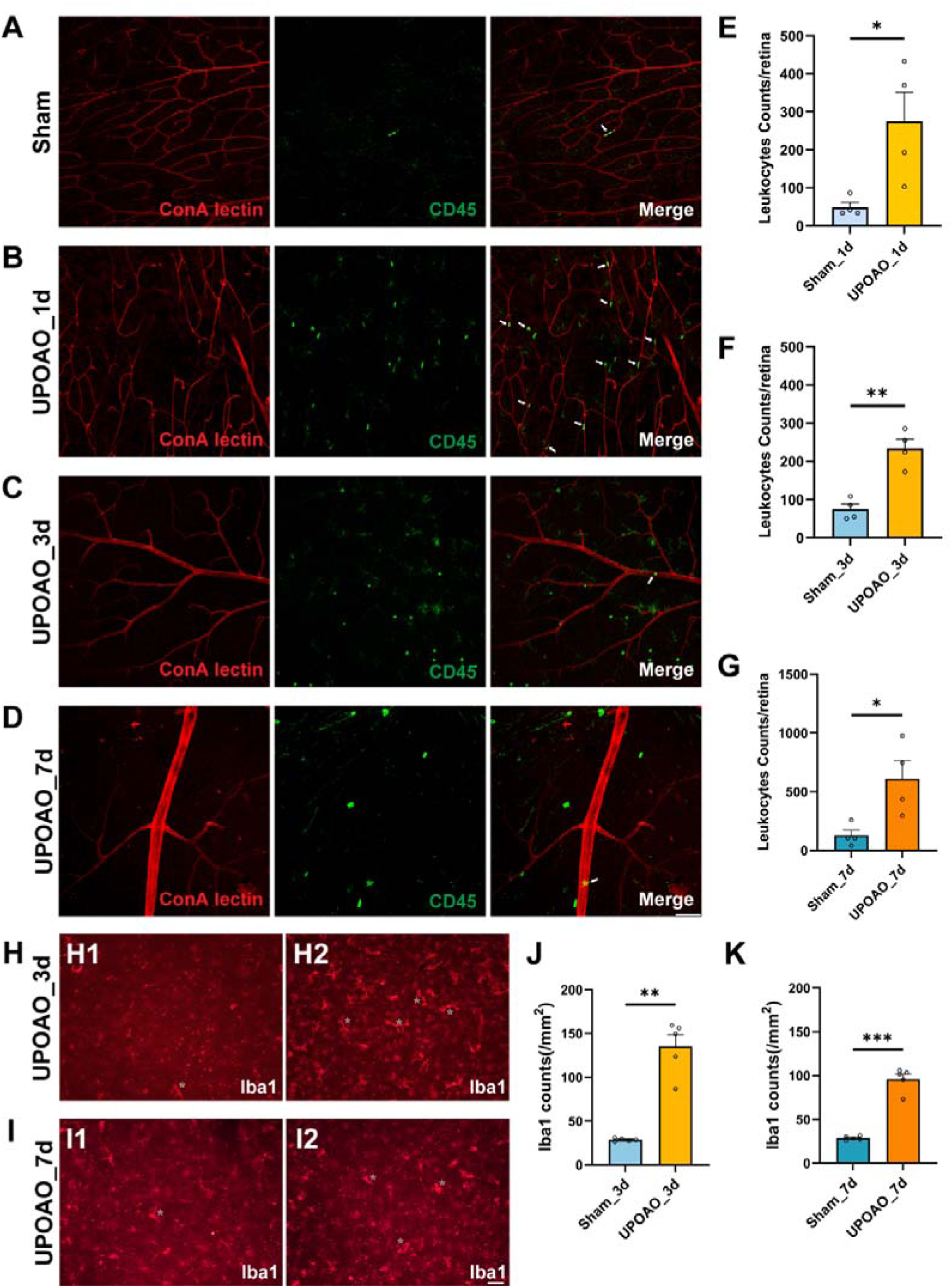
Peripheral Leukocyte Infiltration and Retinal Resident Microglial Activation. Rhodamine-labeled canavalin A was used for immediate cardiac perfusion to visualize blood vessels, followed by CD45 immunofluorescent staining to observe the relationship between blood vessels and CD45^+^ cells in sham (A), 1-day perfusion group (B), 3-days perfusion group (C), and 7-days perfusion group (D). The presence of CD45^+^ cells within blood vessels is indicated by white arrows. Scale bar = 50 μm. CD45^+^ cell counts were performed in whole retinas of 1-day (E), 3-days (F), 7-days (G). The cellular morphology and distribution of microglial cells in the superficial retina were assessed in 3-days (H) and 7-days (I). Activated microglial cells are indicated by white asterisks. Microglial cell counting was conducted in the superficial retina of 3-days (J) and 7-days (K). Data points for CD45^+^ cells were derived from four flat-mounted retinas, and data points for microglial cells were from five flat-mounted retinas. Data were presented as means ± s.e.m, *: p<0.05, **: p<0.01, ***: p<0.001, ****: p<0.0001, t-test. Scale bar = 50 μm.

Microglia, which are known to undergo activation and morphological changes in response to various insults, were examined in the superficial retinas of sham and UPOAO eyes. Immunofluorescence staining for Iba1 was conducted on flat-mounted retinas at 3-days and 7-days post-reperfusion. In sham eyes, Iba1^+^ microglia displayed small somas and elongated dendrites that were evenly distributed within the retina (**Fig. 7H1, I1**). Conversely, UPOAO retinas exhibited numerous activated Iba1^+^ microglia with enlarged somas, shortened dendrites, and an ameboid appearance (**Fig. 7H2, I2**). The number of Iba1^+^ microglia in UPOAO retinas was approximately five times greater than that in sham eyes at 3-days (**Fig. 7J**), with a further increase observed at 7-days (**Fig. 7K**), although this increase was less pronounced than that at the 3-days time point (**Fig. 7J**). These results indicated a potential reduction in immune-mediated inflammation over time.

### 3.7 RNA-seq comparison between UPOAO and extravascular occlusion models

To further study the characteristics of UPOAO, we analyzed the transcriptomes of two extravascular occlusion models: HIOP and UCCAO. The HIOP model induces ischemia through anterior chamber perfusion with normal saline (**Fig. 8A**), while the UCCAO model involves ligation of the unilateral common carotid artery (**Fig. 8H**). RNA-seq analysis of retinas subjected to 60 minutes of ischemia followed by 7-days of reperfusion in the HIOP model revealed enrichment of immune-related pathways, specifically highlighting increased activity in leukocyte-mediated immunity and regulation of immune effector processes (**Fig. 8B**). This finding was consistent with the results of GO analysis, which revealed associations of the overlapping DEGs between the HIOP and UPOAO models with toll-like receptor binding, T-cell receptor binding, and immune-related functions (**Fig. 8C, D**). To further explore these DEGs, we explored the remaining DEGs in the HIOP model in addition to the overlapping genes. GO analysis revealed that the remaining DEGs in the HIOP model were also enriched in immune responses, such as the T-cell receptor complex pathway, adaptive immune response, and B-cell-mediated immunity function (**Fig. 8E**). Remarkably, upon examining the remaining DEGs specific to the UPOAO model, GO analysis revealed DEGs distinctly related to lipid and steroid metabolic processes (**Fig. 8F, G**). This observation indicates that the UPOAO model, which is closely related to RAO, involves not only conventional immune responses but also unique regulation of lipid and steroid metabolism.

**Fig. 8.**
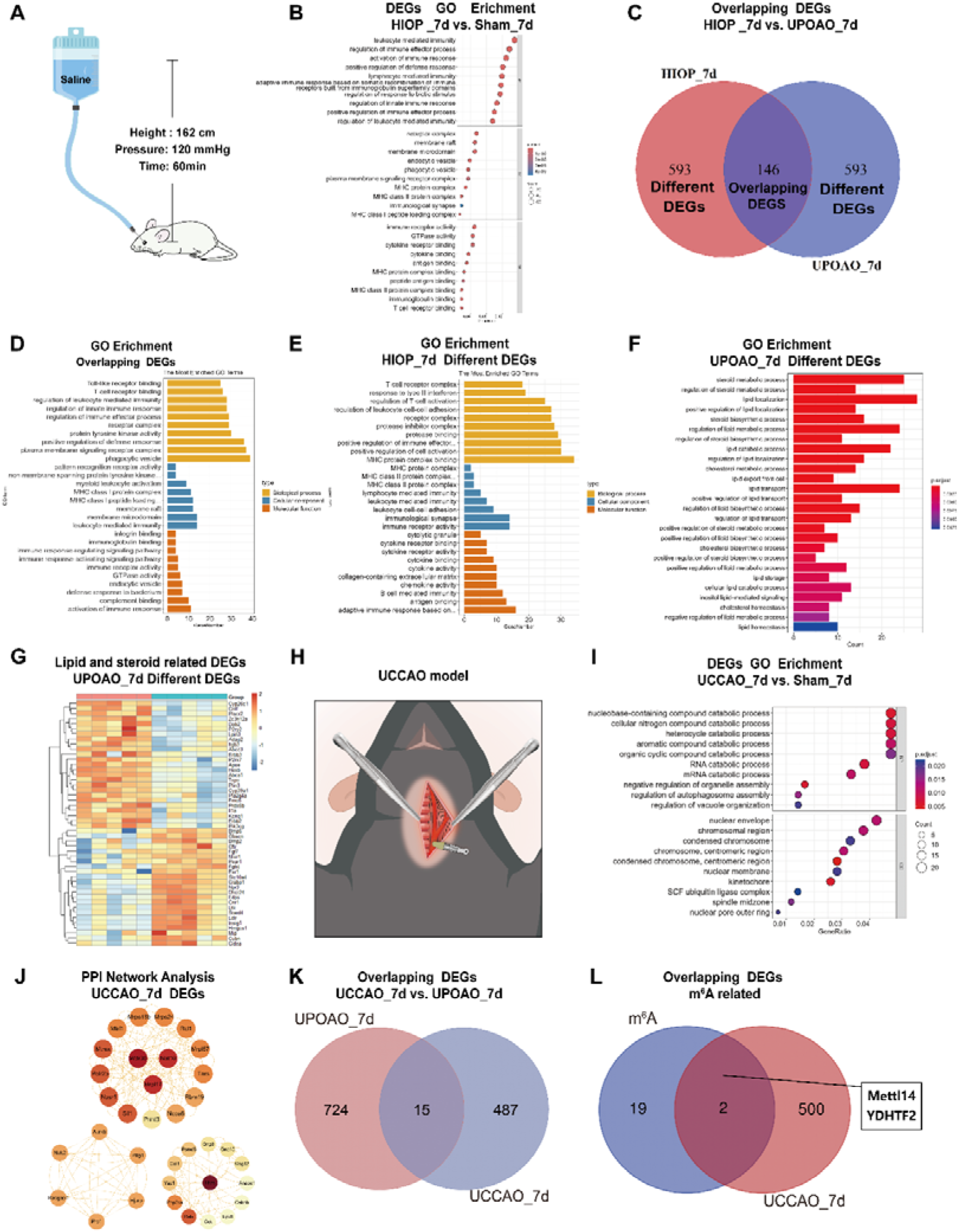
Transcriptomic Results Comparison between UPOAO, HIOP, and UCCAO Models. RNA-seq comparison between UPOAO and extravascular occlusion models: HIOP and UCCAO. (A) Schematic illustration of the HIOP model. (B) GO analysis of differential expression genes (DEGs) in the HIOP model at 7-days perfusion. (C) Venn diagram indicating the overlapping DEGs (146 genes) between HIOP and UPOAO models, as well as the remaining DEGs specific to each group (593 genes in each group). (D) GO analysis of the overlapping DEGs between HIOP and UPOAO models. (E) GO analysis of the remaining DEGs in the HIOP model at 7-days perfusion, excluding the overlapping DEGs. (F) GO analysis of the remaining DEGs, excluding the overlapping DEGs. (G) Heatmap showing the inter-sample distribution of lipid and steroid-related DEGs from the analysis of the remaining DEGs in the UPOAO model. (H) Schematic illustration of the UCCAO model. (I) GO analysis of DEGs in the UCCAO model. (J) Hub genes identified through protein-protein interaction (PPI) network analysis of the DEGs in the UCCAO model. (K) Venn diagram indicating the overlapping DEGs between UCCAO and UPOAO models (15 genes). (L) Venn diagram indicating the presence of two overlapping DEGs between UCCAO DEGs and m6A-related genes.

Similarly, the UCCAO model exhibited associations with negative regulation of B-cell proliferation, lymphocyte activation, and related processes (**Fig. 8I**). Additionally, the hub genes identified through protein‒protein interaction (PPI) analysis are presented (**Fig. 8J**). However, the UCCAO model displayed minimal overlap with the UPOAO model, with only 15 overlapping DEGs identified in the Venn diagram analysis (**Fig. 8K**). Further analysis of the UCCAO DEGs revealed two genes (Mettl14 and YTHDF2) related to m6A modifications (**Fig. 8L**). Although arterial blood flow is occluded in both the UPOAO and UCCAO models, our results revealed that the pathophysiological processes of the two models were particularly different after 7-days of reperfusion.

## 4 Discussion

This study aimed to develop a more appropriate mouse model for investigating the ischemia-reperfusion process in RAO and unravelling the underlying pathophysiological mechanisms. We combined silicone wire embolization with carotid artery ligation to effectively block the blood supply to the retinal artery, closely mimicking the characteristics of the acute interruption of blood supply in RAO patients [31]. In the UPOAO model, a 60-minute ischemia period was sufficient to simulate the injury of major retinal neural cells, such as RGCs, BCs, HCs, and cholinergic amacrine cells, resulting in visual impairment. Moreover, histologic examination demonstrated thinning of the inner layer of the retina, especially the GCL. Time course transcriptome analysis revealed various pathophysiological processes related to immune cell migration, oxidative stress, and immune inflammation during non-reperfusion and reperfusion periods. The resident microglia within the retina and peripheral leukocytes that access the retina were markedly increased during reperfusion periods. Additionally, we compared the transcriptomic signatures of the UPOAO model with those of two commonly used ischemia-reperfusion mouse models (HIOP and UCCAO), highlighting the potential relevance of the UPOAO model to ocular vascular occlusive diseases.

The UPOAO mouse model effectively simulates the characteristic features of RAO observed in patients. RAO patients often exhibit distinctive patterns in dark-adapted ERGs, where b-waves are reduced while a-waves remain stable [32–34]. In our study, we observed almost the same phenomenon in ERG amplitude after 60 minutes of ischemia in the UPOAO model (Table S2). This observation was further confirmed by immunofluorescence staining of bipolar cells and photoreceptor cells. Importantly, a previous study linked the thinning of the inner retina with alterations in ERG in patients with RAO [35]. Our results showed that ischemic damage predominantly affected the inner retinal layers, while the outer layer was almost unaffected (**Table S2**). This distinction is attributed to the reliance of the inner layers on the retinal artery for blood supply, whereas the outer layers are supplied by the choroid. Additionally, histological examination revealed increased retinal NFL and GCL thickness and decreased IPL thickness after 3-days of reperfusion, indicating that ischemia-induced oedema had not completely subsided.

This study comprehensively explored the transcriptomic signature after ischemia and revealed that resident and peripheral immune cells may play major roles in pathological processes. In the non-reperfusion group, the transcriptional changes were primarily involved in the migration of immune cells such as leukocytes and neutrophils from blood vessels into ischemic retinal tissue. This process is closely linked to injury during ischemia and reperfusion, where leukocytes and neutrophils infiltrate neural tissue through the vascular endothelium [36, 37]. Our results revealed an increase in CD45^+^ leukocytes in the retina and retinal microvasculature, suggesting that these leukocytes were recruited from the bloodstream to the damaged retinal tissue following reperfusion. During leukocyte access to the retina, chemokines, chemokine receptors, adhesion molecules and cytoskeletal components play important roles in regulating endothelial permeability and facilitating leukocyte adhesion to endothelial cells [38, 39]. Our findings of related gene expression are consistent with this mechanism.

During the 3-days reperfusion period, we observed that an increased number of DEGs were significantly involved in the critical pathological response to oxidative stress in the context of IR tissues. The generation of reactive oxygen species (ROS) in mitochondria and subsequent oxidative stress are widely recognized as major causes of retinal cell damage induced by IR [40, 41]. This finding suggested that mitochondrial function, which is involved in oxidative stress processes, may play an important role in the pathophysiology of IRI during the early reperfusion stage. Oxidative stress can influence the permeability of the inner blood retina barrier (iBRB), allowing leukocytes to access retinal tissues [42]. The infiltration and activation of immune cells are recognized as the underlying causes of many retinal diseases, including ischemic ophthalmia [42, 43]. In the UPOAO model, the increase in CD45^+^ leukocytes and Iba1^+^ microglia in the retina after 3-days of reperfusion supports this viewpoint.

In the later stage of reperfusion (7-days), the DEGs were mainly enriched in immune regulation and inflammation. With more upregulated DEGs than downregulated ones, we hypothesize that the activation of the immune inflammatory response contributes to further IRI in retinal tissue. Previous studies have shown that the retina can elicit immunological responses during ischemia-reperfusion injury and that immune inflammation is an important phenomenon in the progression of this injury [30]. The excessive ROS generated by mitochondria in RGCs during the activation of inflammatory responses can damage cell structure and visual function [44]. Microglia, considered the primary resident immune cells, contribute to inflammatory responses and consequent neural damage [45]. Infiltrating leukocytes can activate resident microglia, forming a feedback loop that exacerbates inflammation [46]. Complex interactions between immune cells and retinal neurons after RIRI have been reported [47]. Interestingly, in our UPOAO model, the death of major retinal neural cells, such as RGCs, BCs, and HCs, was correlated with an increase in infiltrating leukocytes and resident microglia. Therefore, we considered that the immune response and neuroinflammation observed after 7-days of reperfusion may result from the cumulative effects of acute oxidative stress and the responses of resident and peripheral immune cells (**Fig. 9, Table S2**). In summary, we described the pathological processes at different time points and highlighted the important role of resident and peripheral immune cell responses in the UPOAO model. This finding offers valuable insights for subsequent screening of pathogenic genes and potential immunotherapy approaches.

**Fig. 9.**
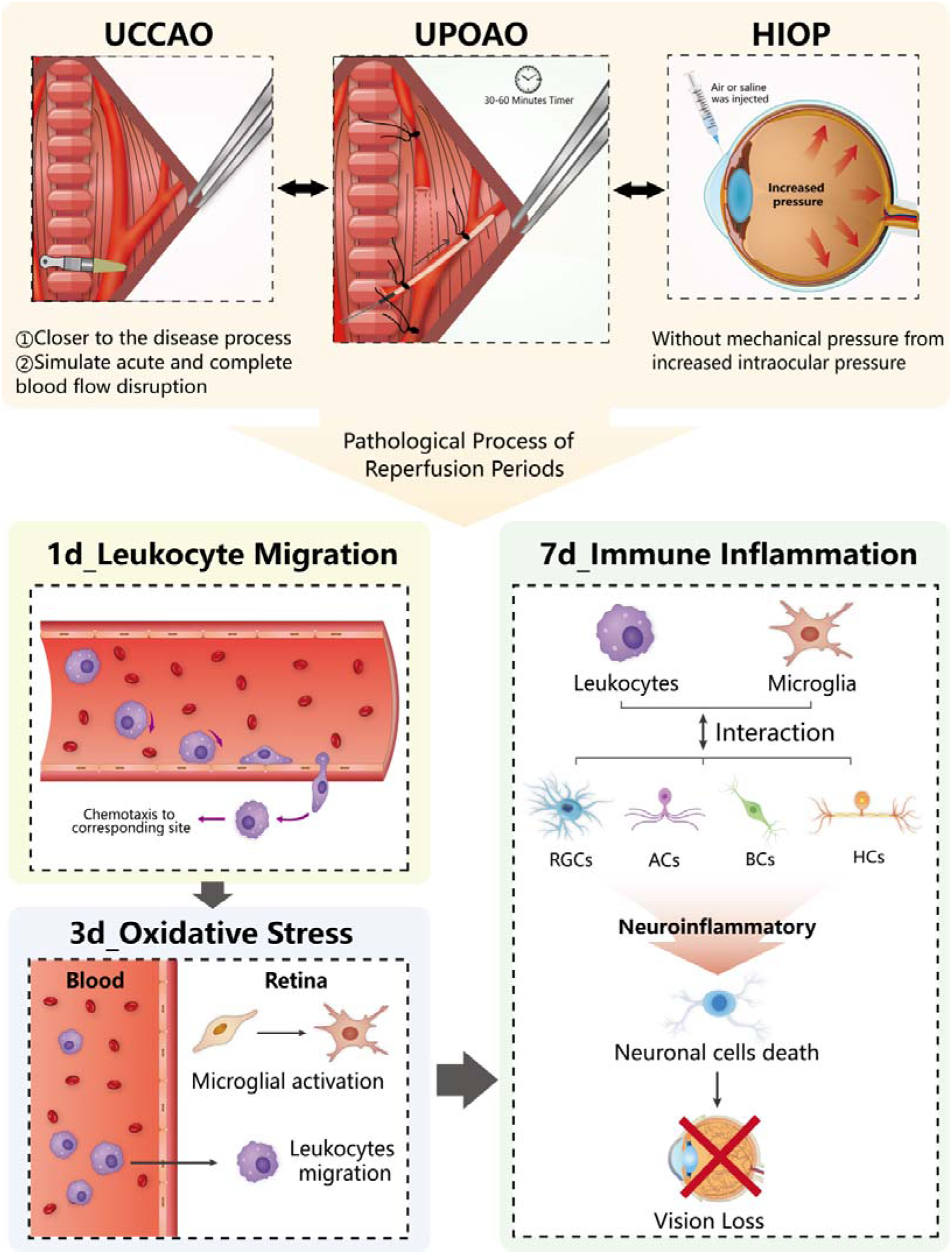
The Characteristic Features of UPOAO Model during Ischemia-Reperfusion Periods.

Compared with the HIOP and UCCAO models, the UPOAO model is a more appropriate choice for studying retinal IRI in RAO. The HIOP model, widely used for investigating the pathogenesis of APACG, exhibits retinal degeneration features similar to those of the UPOAO model during both reperfusion periods [48]. In our UPOAO model, histologic and immunohistochemical analysis demonstrated that 60 minutes of ischemia followed by 3-days or 7-days of reperfusion can cause irreversible damage to retinal structure and visual function. Specifically, we observed distinct alterations, such as thinning of the inner retina, substantial RGCs apoptosis, and decreased b-wave amplitude. Multiple studies on the HIOP model have reported consistent results [45, 49, 50]. However, some typical phenotypes in the HIOP model, such as thinning of the outer nuclear layer and the entire neuroretina and decreased a-wave amplitude, were not observed in the UPOAO model. Moreover, our transcriptome sequencing revealed a specific subset of DEGs unique to the UPOAO model, including Apoe, Abca1, Ldlr, Cyp39a1, and Bmp6. Notably, these DEGs were significantly enriched in pathways associated with lipid and steroid synthesis. Steroid and lipid metabolism homeostasis is disrupted under conditions of oxidative stress and inflammation [51]. In the UPOAO model, the observed lipid and steroid biological processes may result from the cumulative pathological responses triggered by IRI, as observed in the HIOP model. In the UCCAO model, RNA-seq revealed that DEGs were enriched in two main pathways: 1) epigenetic modification-related pathways, including nucleobase-containing compound catabolic process, RNA catabolic process, mRNA catabolic process and N6-methyladenosine (m6A) modification; and 2) cell death pathways, including regulation of autophagosome assembly, negative regulation of neuron death, and negative regulation of the neuronal apoptotic process. Epigenetic mechanisms may play a key role in the pathophysiology of ocular disease [52]. m6A, one of the most common RNA modifications, has been reported to regulate cell death processes, including apoptosis and autophagy, in the pathological process of IRI [53]. These results suggest that epigenetic mechanisms may significantly influence cell death during the 7-days reperfusion period in the UCCAO model. Lee et al. evaluated the characteristics of UCCAO without reperfusion using visual, histological and immunohistochemical approaches. Their results revealed delayed perfusion of the ipsilateral retina, thinning of the inner retinal layer 10 weeks after surgery, and a dramatic decrease in the amplitudes of b-waves on day 14 after UCCAO [15, 54]. This finding suggested that UCCAO primarily represents a model of retinal hypoperfusion injury and may not effectively reflect acute ischemic-induced structural and functional damage in RAO (**Fig. 9**). These results led us to consider the UPOAO model an effective experimental model for studying the pathological processes underlying acute ischemia and IRI.

Our research has certain limitations that should be acknowledged. First, our study focused on the changes that occurred within 60 minutes of ischemia and within the first 7-days of reperfusion in the UPOAO model. Further exploration is needed to understand the changes induced by longer reperfusion periods. Additionally, we proposed possible pathological mechanisms, including iBRB damage, oxidative stress and immune inflammation, mainly based on the enrichment results of DEGs at three reperfusion time points. More molecular experimental validation is required to substantiate these proposed mechanisms.

In summary, by thoroughly exploring the injury to major retinal neural cells, visual impairment and pathophysiological changes in the UPOAO model, we confirmed that this model can effectively simulate the acute ischemia-reperfusion processes in RAO. It serves as an ideal mouse model for investigating the underlying pathological mechanisms of ischemia and reperfusion. Furthermore, our UPOAO model holds great promise as a novel model for studies on pathogenic genes and potential therapeutic interventions for RAO.

## Acknowledgements

The authors would like to express their gratitude to all participants involved in the experiments. The authors also wish to acknowledge the Eye Institute of Renmin Hospital of Wuhan University for providing us with the experimental facilities and environment. Special thanks are extended to Professor Shenqi Zhang for the technical support in the surgical procedure of UPOAO model.

## Availability of data and materials

All data are available from the corresponding author upon reasonable request.

## Authors’ contributions

Xiao X and Yang AH conceived the project and supervised, guided the research. Li Y, Wang YD and Feng JQ wrote the draft, Feng JQ, Wang CS, Xie H and Li ZY revised the manuscript. Wang YD and Li Y constructed the animal models and obtained the data. Wang YD performed the immunofluorescence staining, HE staining. Li Y and Feng JQ performed the ERG, analyzed the RNA-seq data, and created the visualizations. Wan YW and Wang YD performed the FFA and OCT imaging and Wan YW analyzed the data. Lv BY and Wang YD took pictures and videos of the UPOAO model, and Feng JQ made schematic diagrams of modeling. Li YM, Xie H, Chen T, Wang FX provided support for the technical methods and contributed to the discussion, and reviewed the manuscript. All authors critically contributed to the manuscript, and have read and approved the final manuscript.

## Funding

This work was supported by the National Nature Science Foundation of China (grant number: 82371079), National Key R&D Program of China (2023YFC2308404), Fundamental Research Funds for the Central Universities (2042023gf0013), Key research and development project of Hubei Province (grant number: 2022BCA009).

## Ethics approval and consent to participate

All animal experiments were performed following the ethical guidelines outlined in the ARVO Statement for the Use of Animals in Ophthalmic and Vision Research, with the approval of the Experimental Animal Ethics Committee of Renmin Hospital of Wuhan University (approval number: WDRM-20220305A).

## Supplementary data

**Fig. S1.**
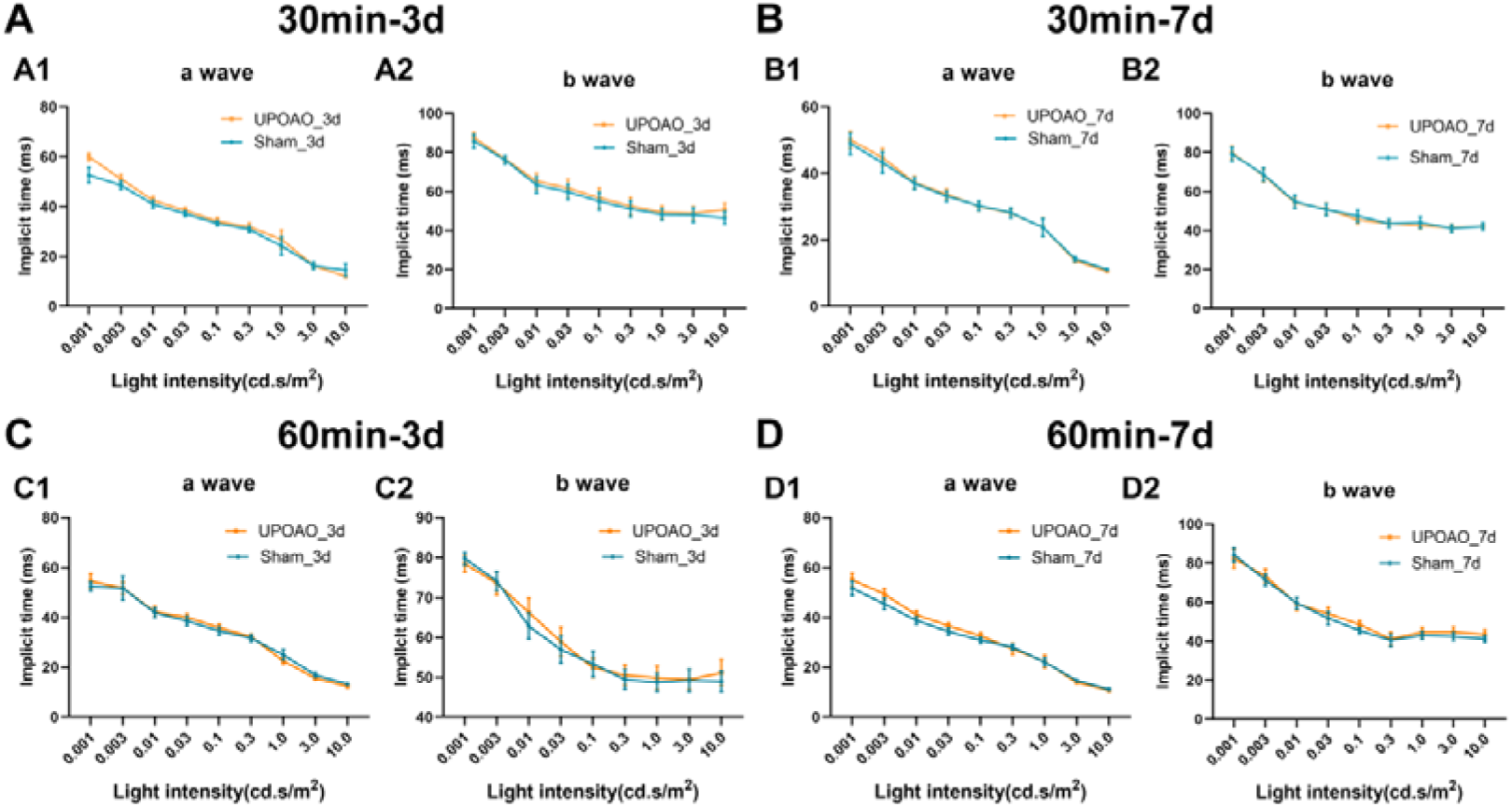
Response Times of a-Waves and b-Waves in ERG at Different Light Intensities. (A, B) Response times of a-waves and b-waves were measured under different light intensities at 3-days and 7-days post-UPOAO in the 30-minutes ischemia group. n = 5. (C, D) Response times of a-waves and b-waves were measured under different light intensities at 3-days and 7-days post-UPOAO in the 60-minute ischemia group. n = 7 at 3-days; n = 8 at 7-days. Data were presented as means ± s.e.m, *: p<0.05, **: p<0.01, ***: p<0.001, ****: p<0.0001, two-way ANOVA test.

**Fig. S2.**
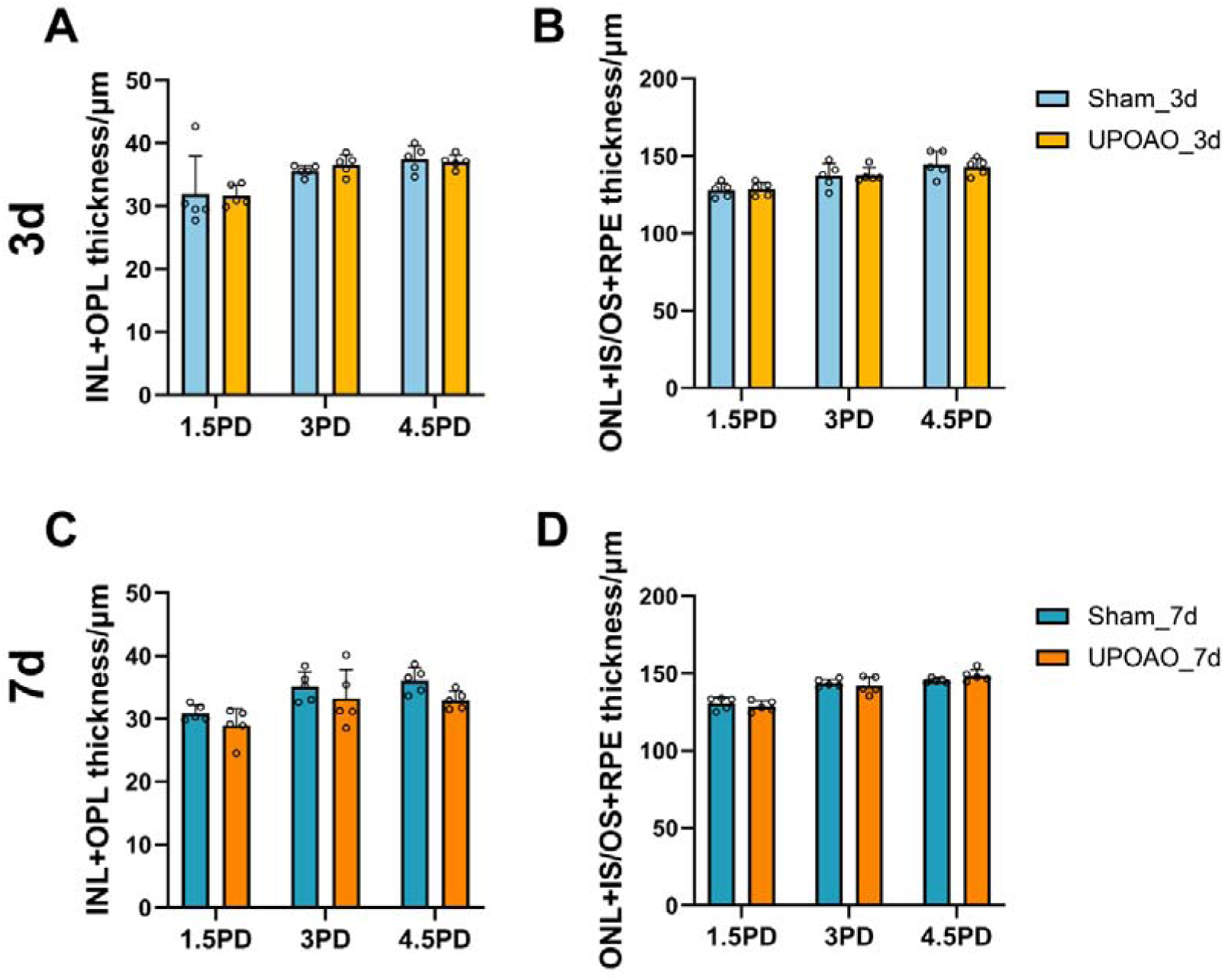
Quantification of INL+OPL Thickness and ONL+IS/OS+RPE Thickness in OCT during 3-days and 7-days Reperfusion of UPOAO Animals. The INL+OPL thickness and the ONL+IS/OS+RPE thickness were measured and compared at distances of 1.5 PD, 3.0 PD and 4.5 PD from the optic disc using OCT. (A and B) Quantification of INL+OPL layer thickness (A) and ONL+IS/OS+RPE layer thickness (B) at 3-days. n = 5. No significant differences were observed. (C and D) Quantification of INL+OPL layer thickness (C) and ONL+IS/OS+RPE layer thickness (D) at 7-days. n = 5. No significant differences were observed. Data were presented as means ± s.e.m, *: p<0.05, **: p<0.01, ***: p<0.001, ****: p<0.0001, two-way ANOVA test.

**Fig. S3.**
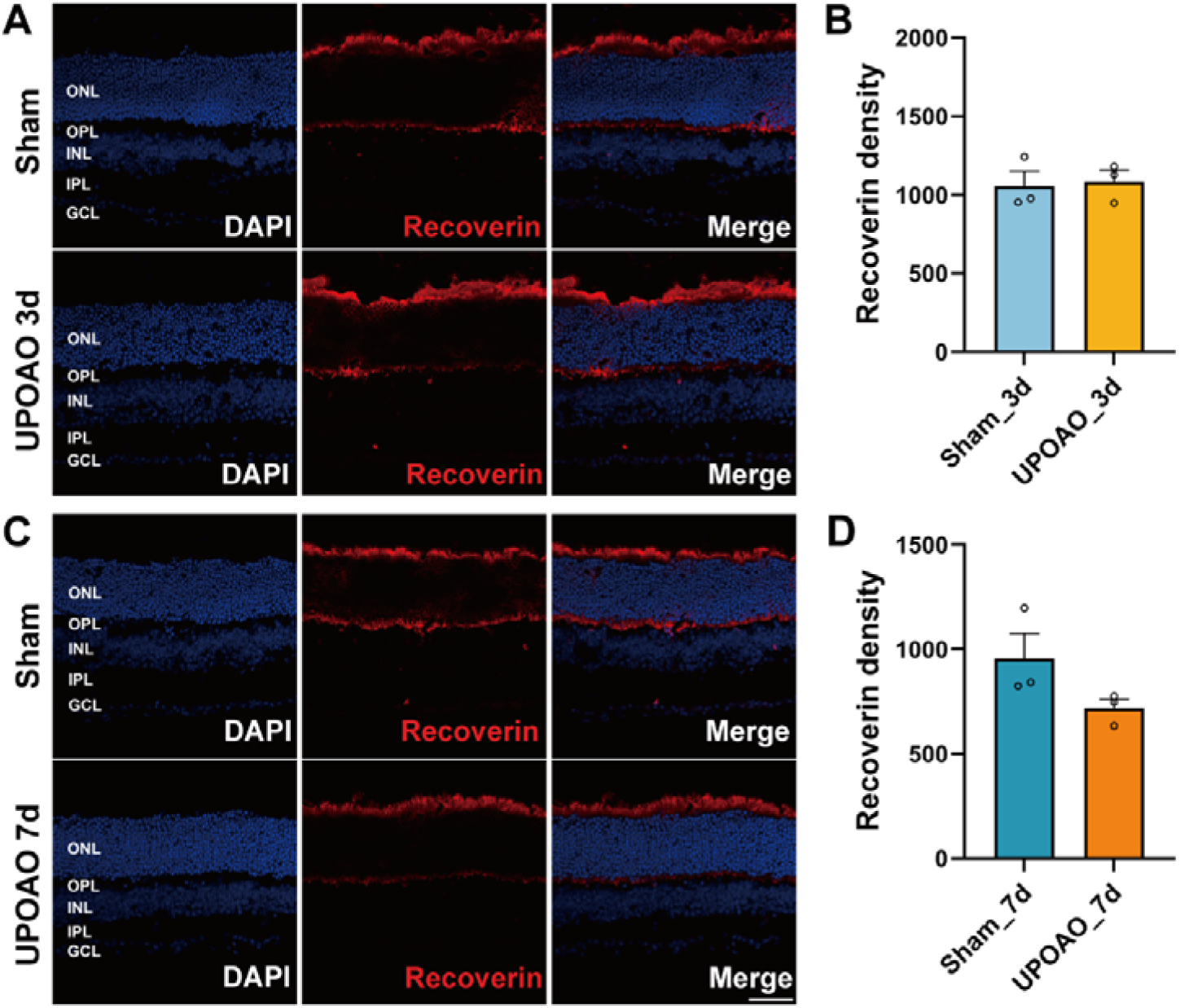
Changes in Photoreceptor Cells in UPOAO. (A) Representative images of mouse retina co-stained with DAPI and Recoverin at 3-days. (B) Representative images of mouse retina co-stained with DAPI and Recoverin at 7-days. n = 3. Data were presented as means ± s.e.m, *: p<0.05, **: p<0.01, ***: p<0.001, ****: p<0.0001, t-test. Scale bar = 50 μm.

**Fig. S4.**
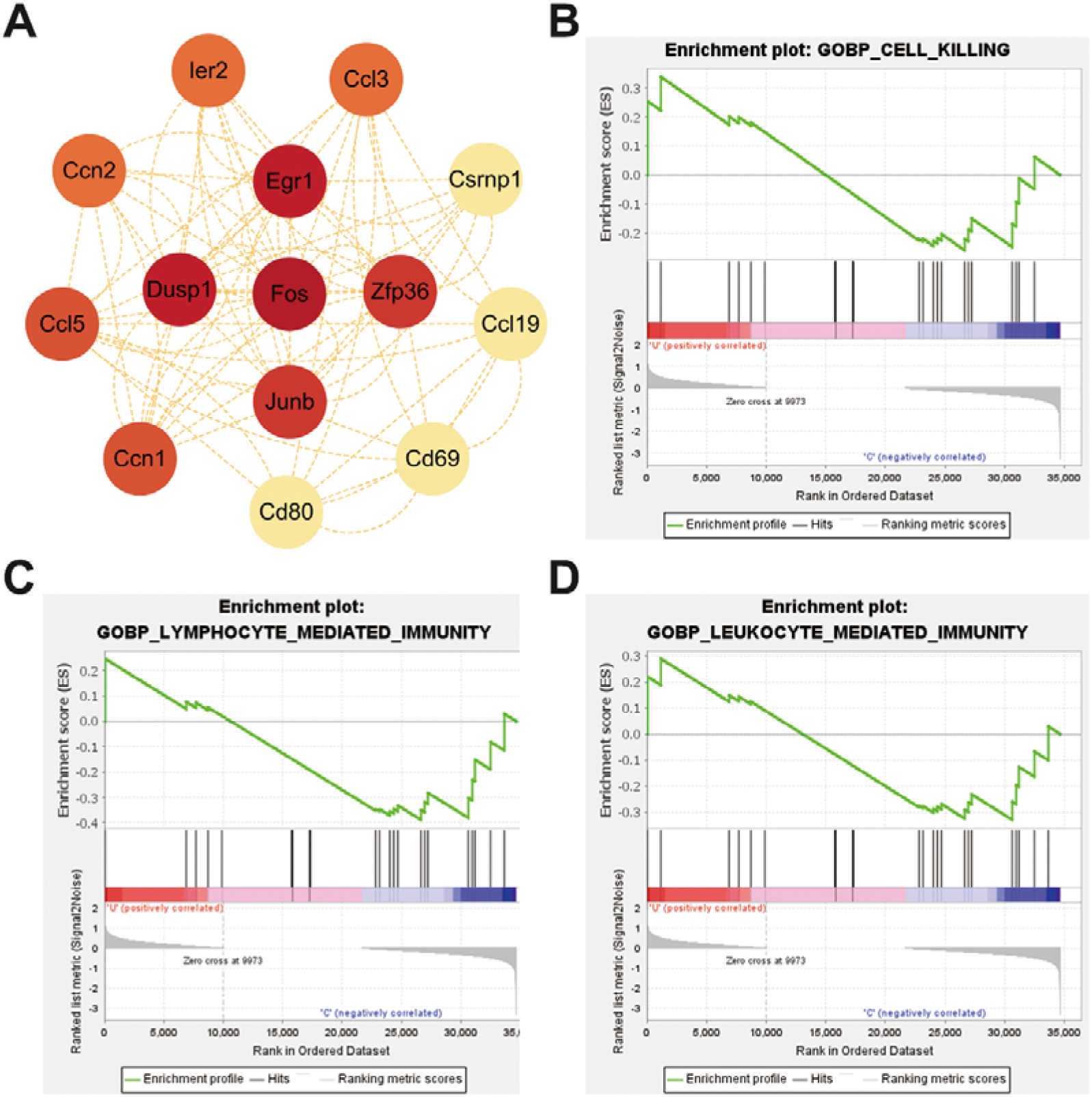
Hub Genes in PPI Analysis and Gene Set Enrichment Analysis (GSEA) during the Non-Reperfusion Stage in UPOAO. (A) Hub genes identified through PPI analysis of DEGs in the non-reperfusion group. (B-D) GSEA analysis of the pathway of the cell killing (B), lymphocyte mediated-immunity (C), and leukocyte mediated-immunity (D) in the non-reperfusion group.

**Fig. S5.**
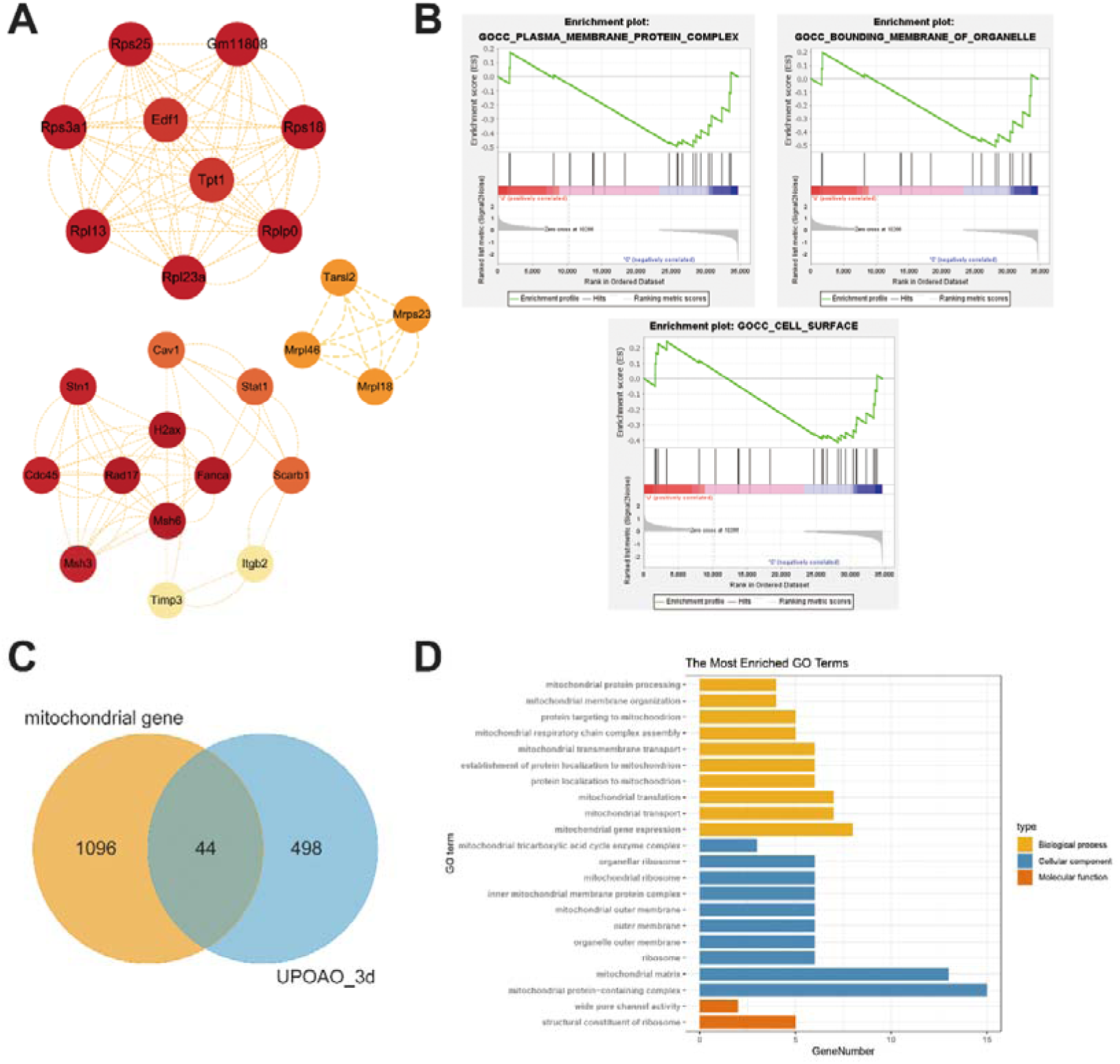
Hub Genes, GSEA Analysis at 3-days Reperfusion in UPOAO, and Relation with Mitochondrial Genes. (A) Hub genes identified through PPI analysis of DEGs in the 3-days reperfusion group. (B) GSEA analysis of the pathways related to plasma membrane protein complex (top left), bounding membrane of organelle (top right), and cell surface (bottom). (C) Venn diagram showing the overlapping genes between mitochondrial genes and DEGs at 3-days reperfusion (44 genes). (D) GO analysis of the 44 overlapping DEGs.

**Fig. S6.**
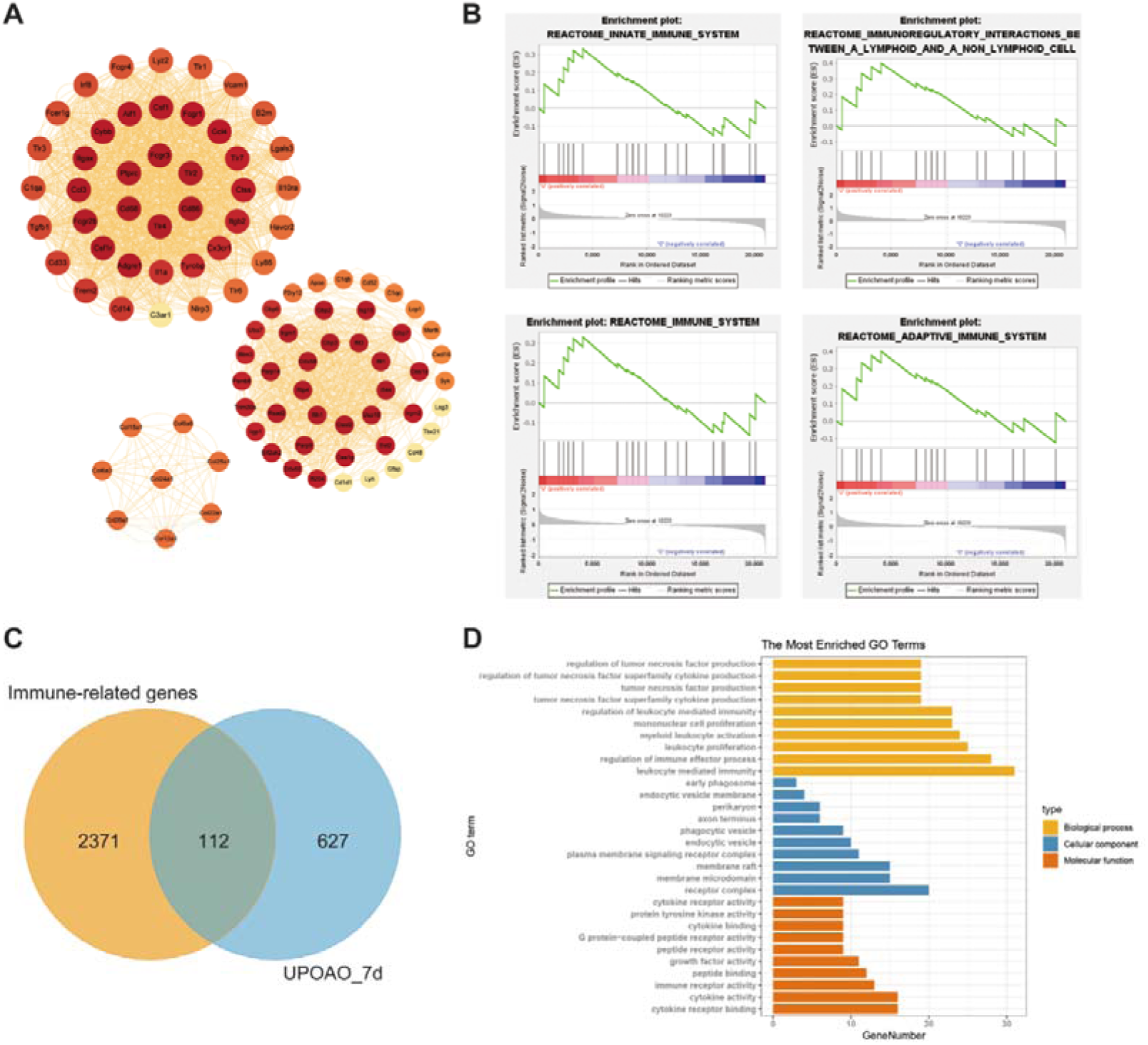
Hub Genes, GSEA Analysis at 7-days Reperfusion in UPOAO, and Relation with Immune Genes. (A) Hub genes identified through PPI analysis of DEGs in the 7-days reperfusion group. (B) GSEA analysis of the pathways related to the reactome innate immune system (top left), reactome immunoregulatory interactions between a lymphoid and a non-lymphoid cell (top right), reactome immune system (bottom left), and reactome adaptive immune system (bottom right). (C) Venn diagram showing the overlapping genes between immune genes and DEGs at 7-days reperfusion (112 genes). (D) GO analysis of the 112 overlapping DEGs.

**Fig. S7.**
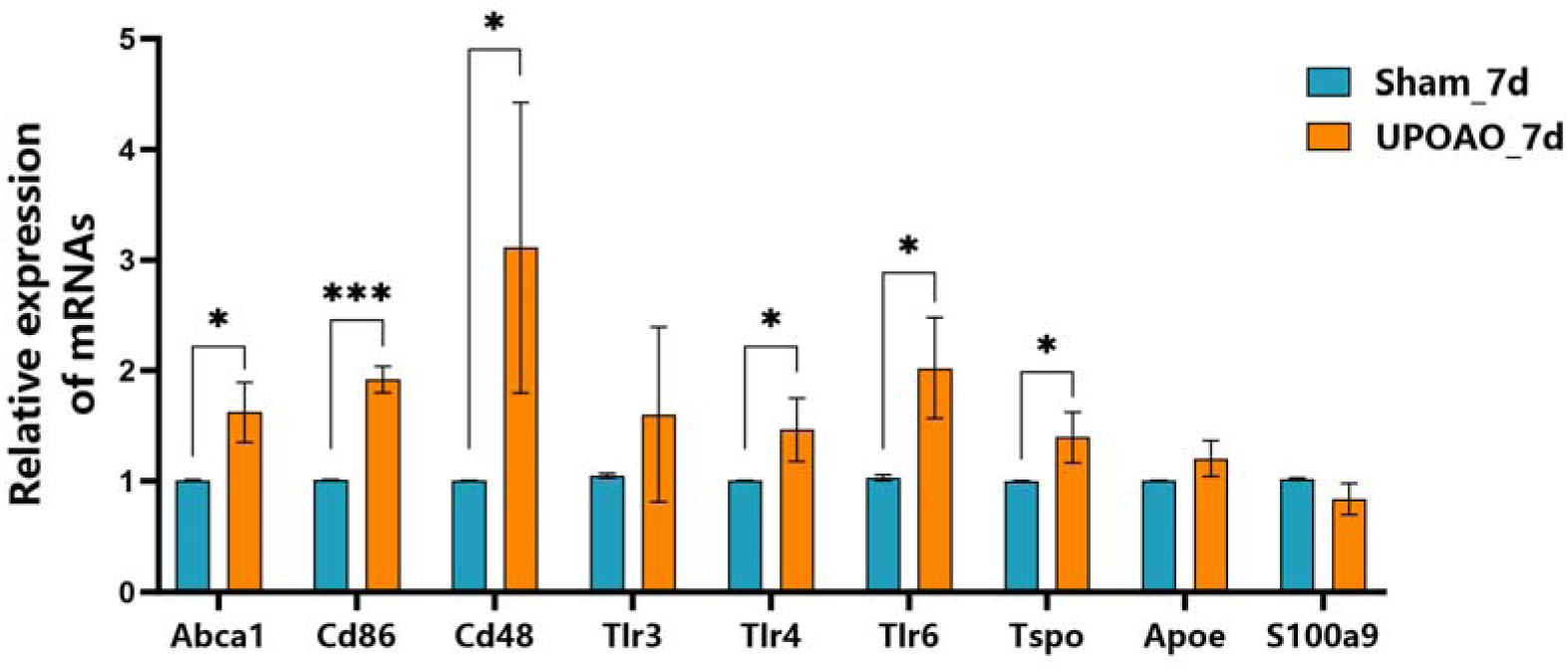
Upregulation of Immune Inflammation-Related Gene Expression in the 7-days Reperfusion Group. The genes Abca1, Cd86, Cd48, Tlr4, Tlr6, and Tspo were significantly upregulated in the 7-days reperfusion group, while other immune-inflammatory and chemotaxis-related genes showed no significant differences. The data points were from the retina of four animals. Data were presented as means ± s.e.m, *: p<0.05, **: p<0.01, ***: p<0.001, ****: p<0.0001, Paired t-test.

**Fig. S8.**
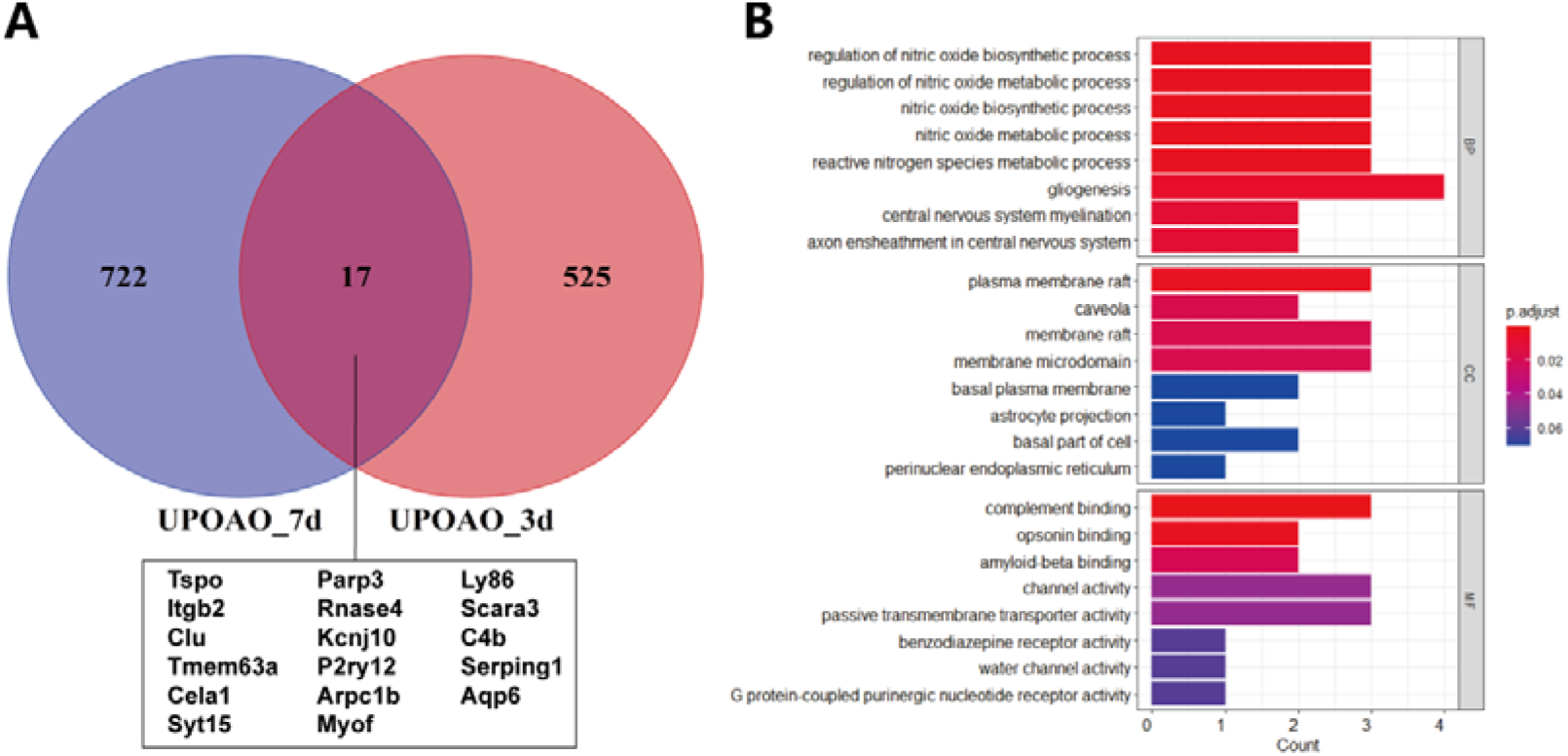
Co-Expressed Genes during 3-days and 7-days Reperfusion. (A) Venn diagram showing the overlap of DEGs between the 3-days and 7-days reperfusion groups, along with a list of the 17 overlapping DEGs. (B) GO analysis of the 17 overlapping DEGs.

**Table. S1.**
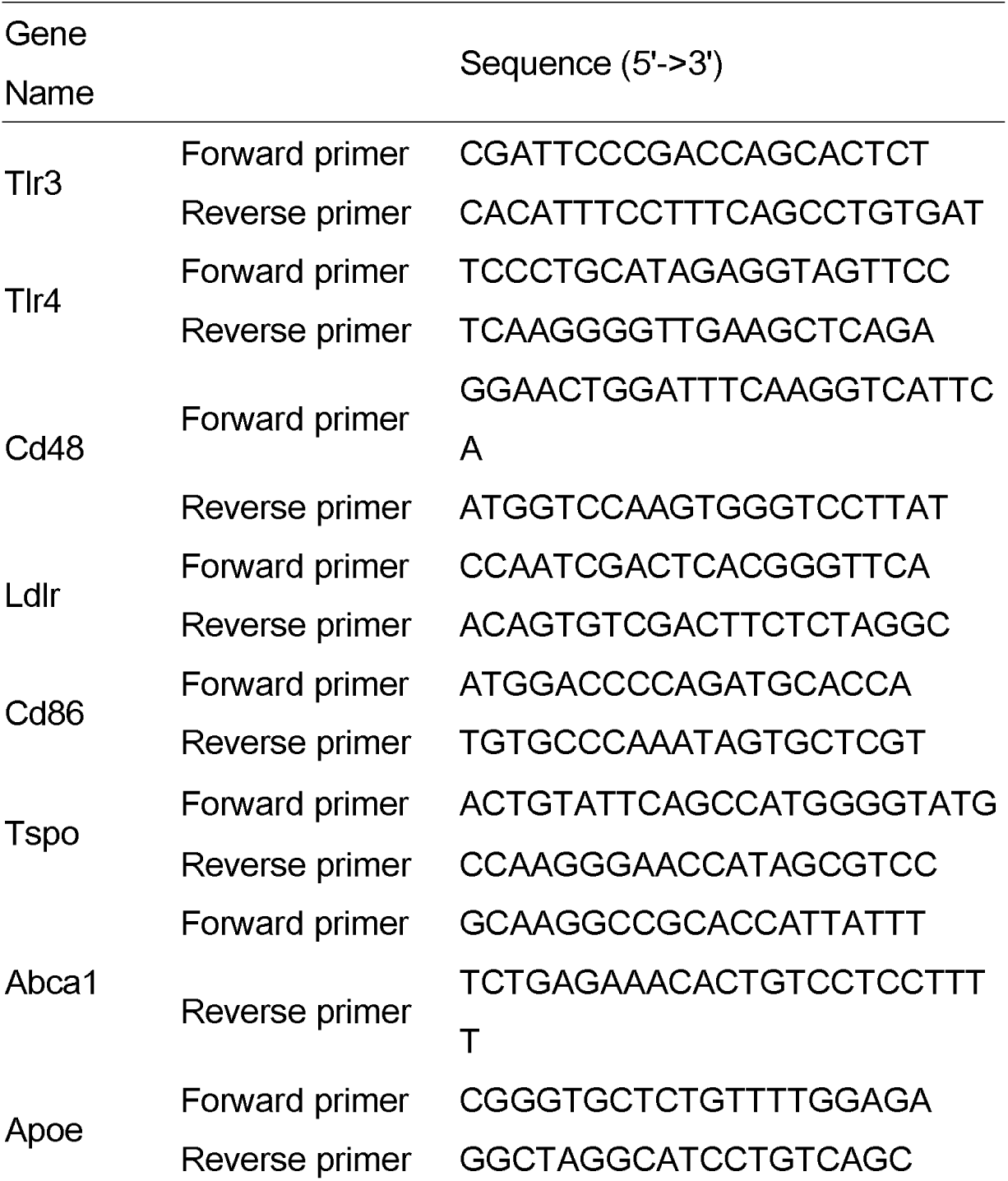
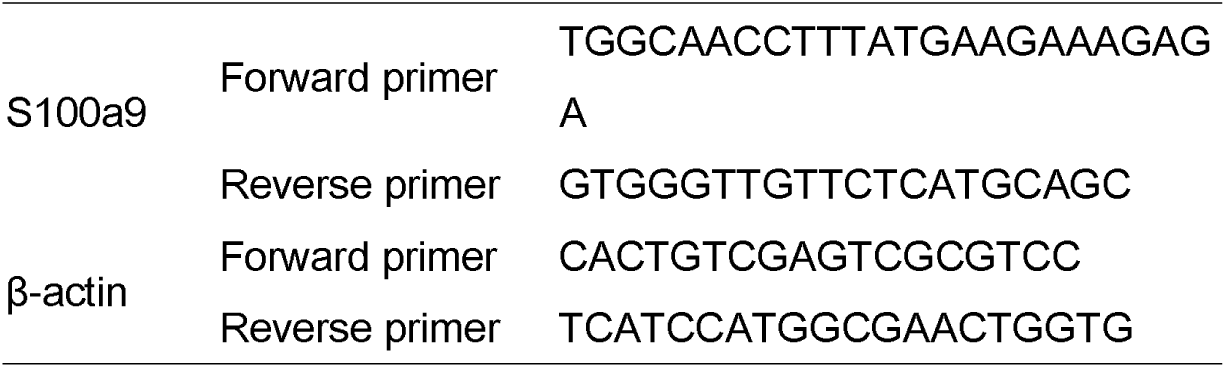
Primers Used in This Study.

**Table. S2.**
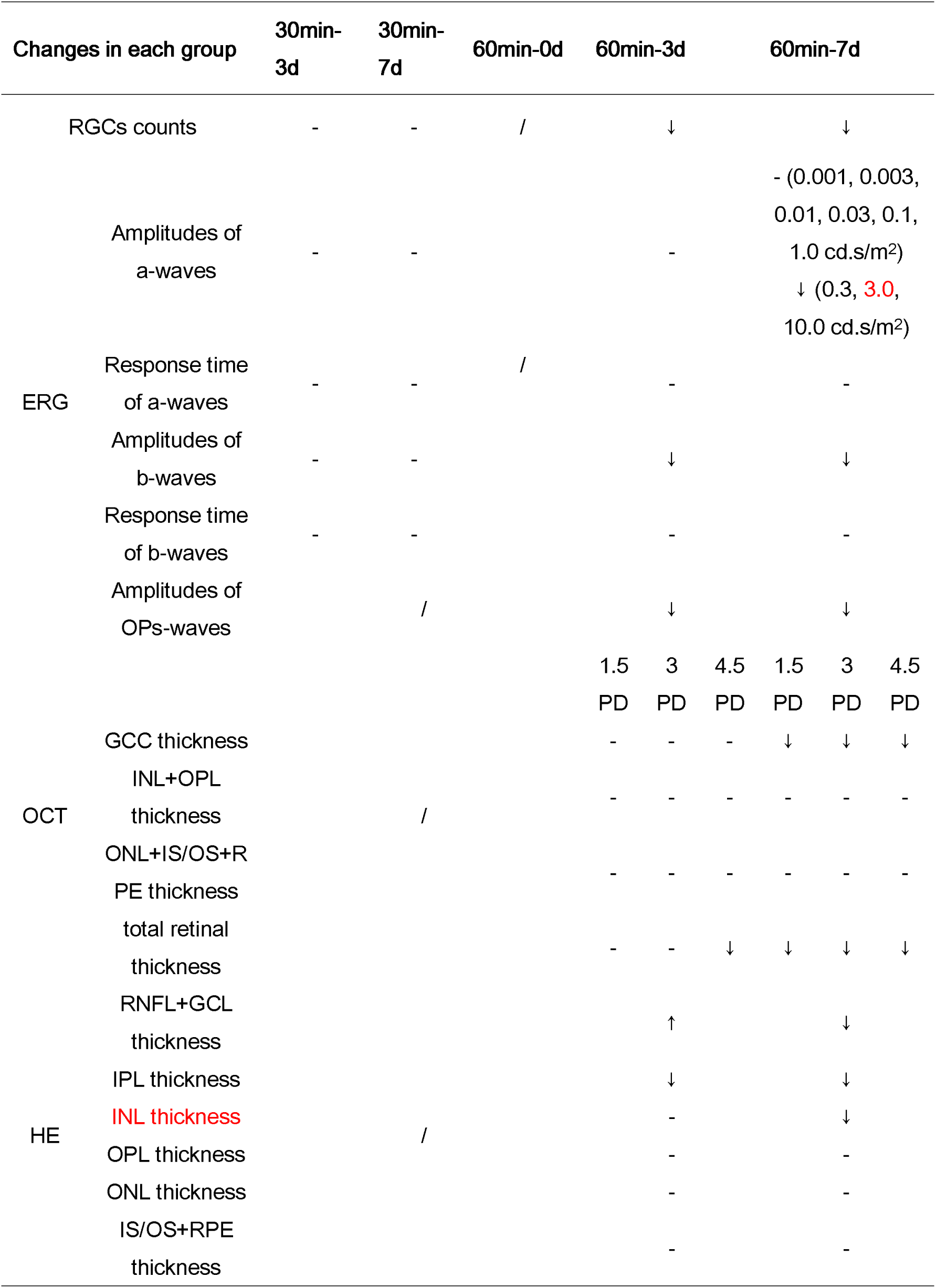
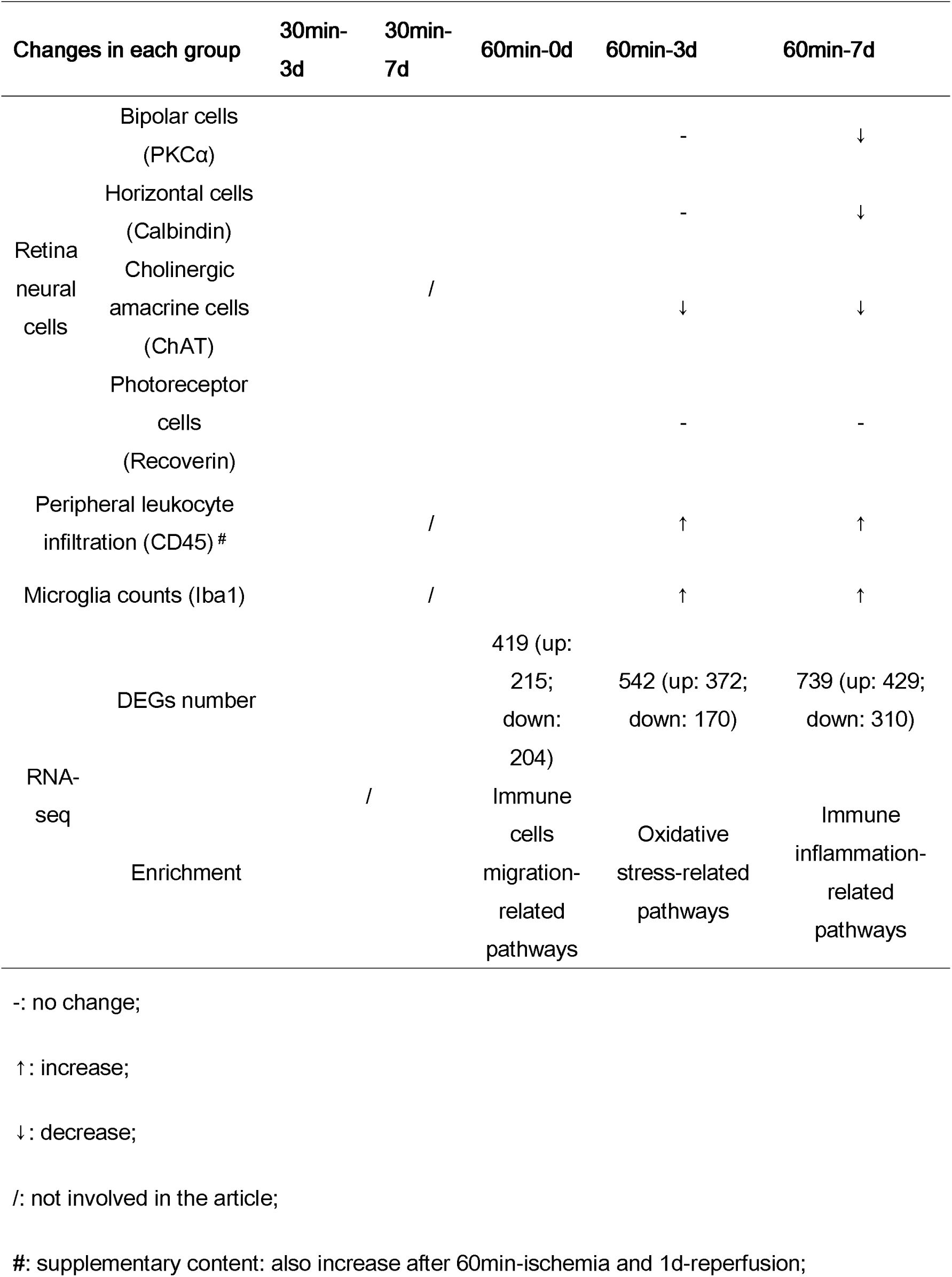
Summary Table: Time Course of all the Morphological, Functional, Cellular, and Transcriptome Changes in the UPOAO Model.

**Video 1. Blunt Dissection of Cervical Vessels.** The neck was exposed, and the cervical arteries were separated using blunt dissection techniques.

**Video 2. Insertion of Silicone Wire Embolus and Induction of Ischemia.** The silicone wire embolus was inserted into the PPA, initiating the ischemic condition. PPA: the pterygopalatine artery.

**Video 3. Extraction of Silicone Wire Embolus and Reperfusion.** The silicone wire embolus was removed, allowing reperfusion to occur. The mouse’s wound was sutured, and standard feeding procedures were followed.

## Notes

### Competing Interest Statement

The authors have declared no competing interest.

### Summary of Updates

Methodology Section Updated: Detailed methodology for RGC quantification was added to the manuscript. Figure 3 Corrected: The duplicated image in Fig. 3E-b wave was replaced with the correct one. Figure 4 Revised: Retinal layer thickness quantifications in HE-stained sections were corrected, including recalibration of the microscope scale and updating thickness measurements. Summary Table Corrected: Typos in the summary table were corrected, including the adjustment of 'Amplitudes of a-wave' and 'INL thickness'.

