## Supplementary figures and images for "Silicone Wire Embolization-induced Acute Retinal Artery Ischemia and Reperfusion Model in Mouse: Gene Expression Provides Insight into Pathological Processes"

### Supplemental Figure 1

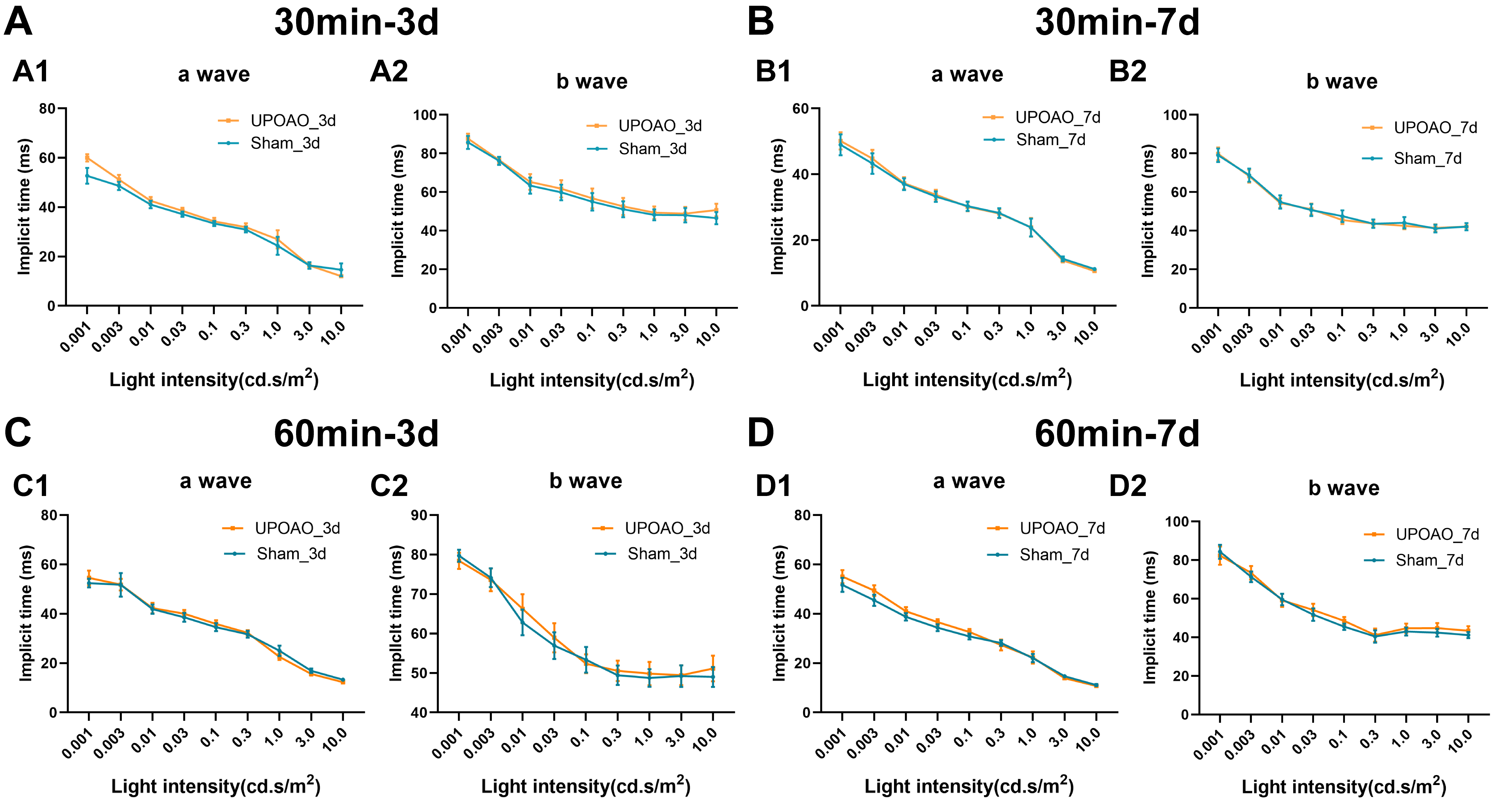

### Supplemental Figure 2

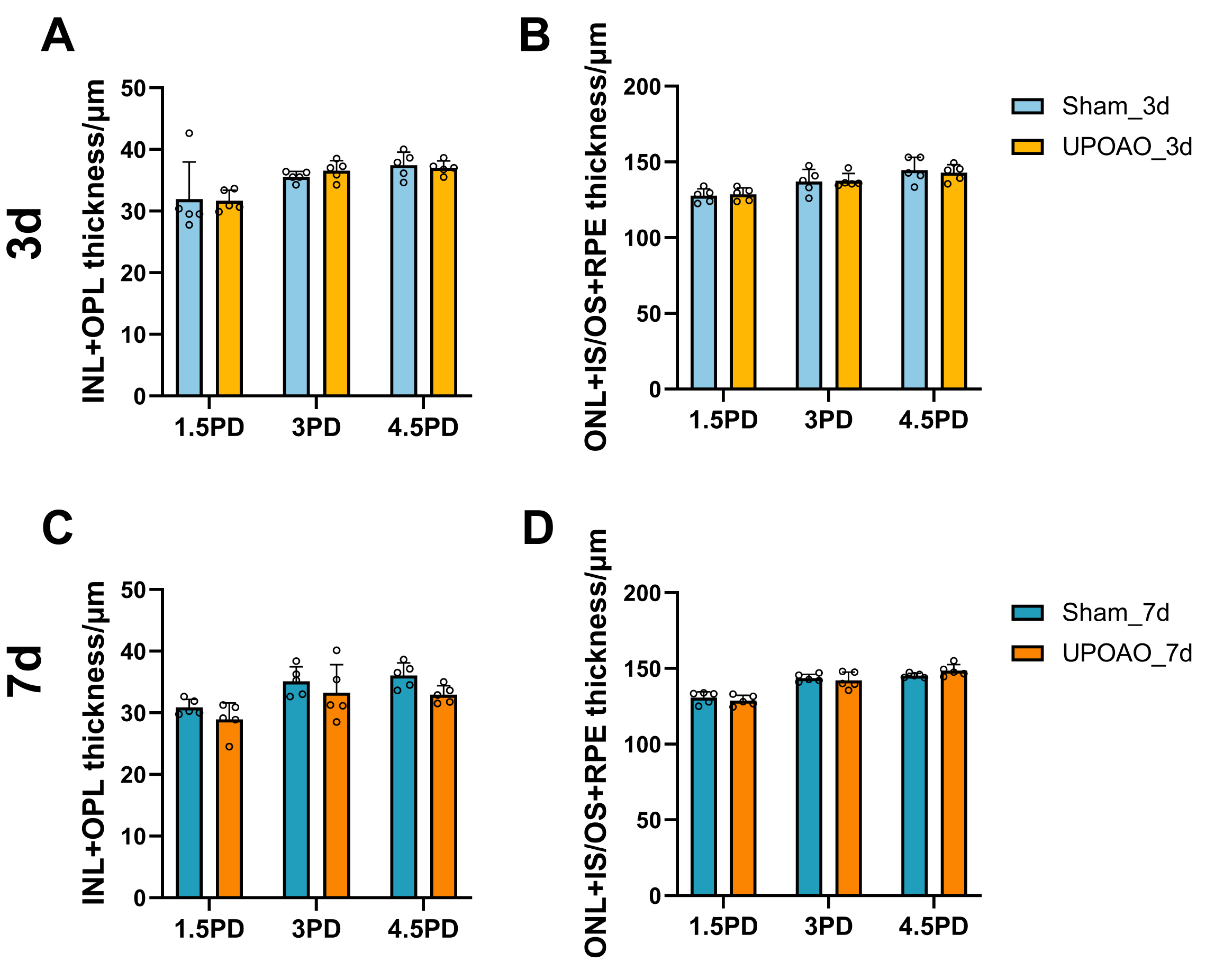

### Supplemental Figure 3

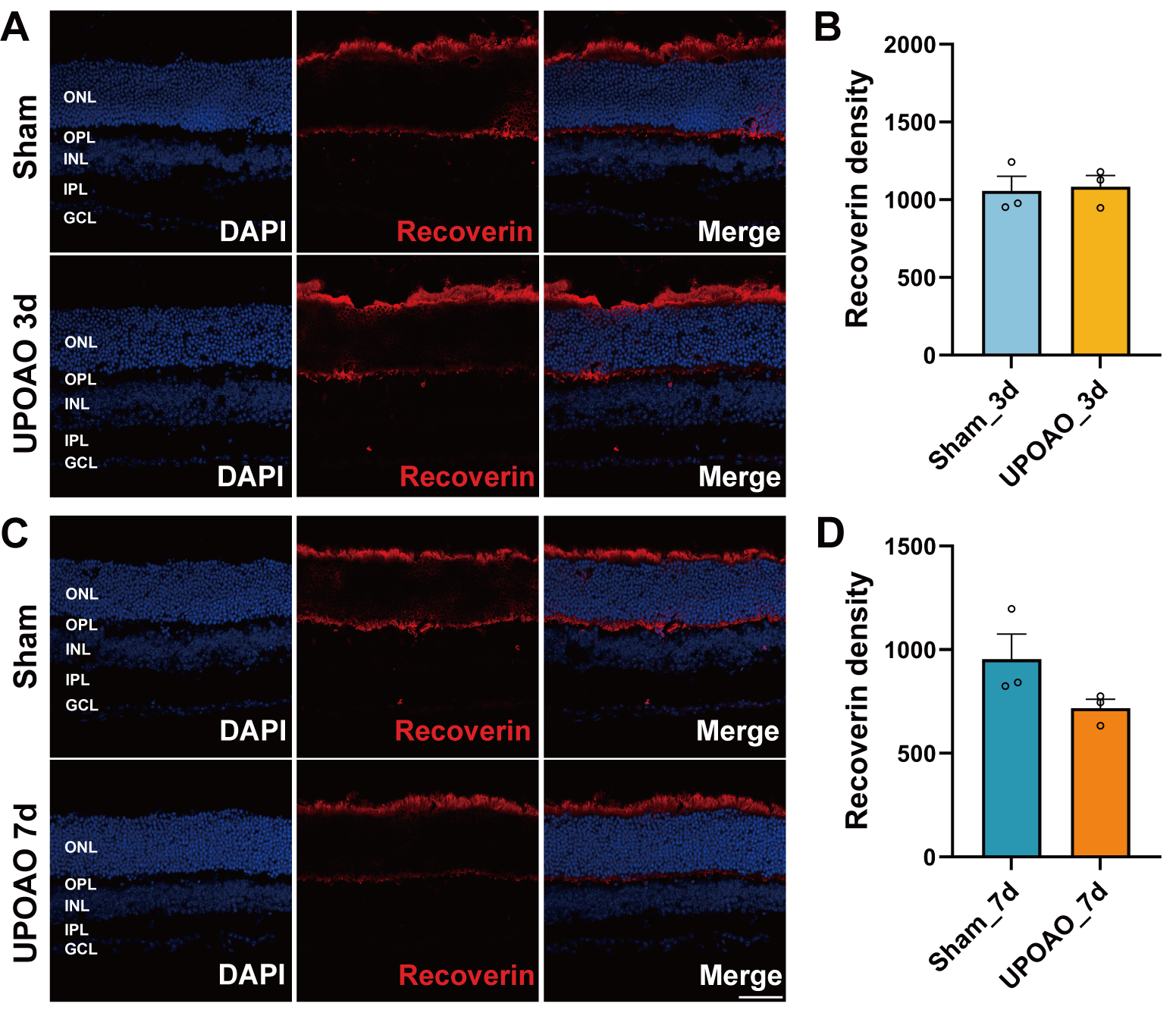

### Supplemental Figure 4

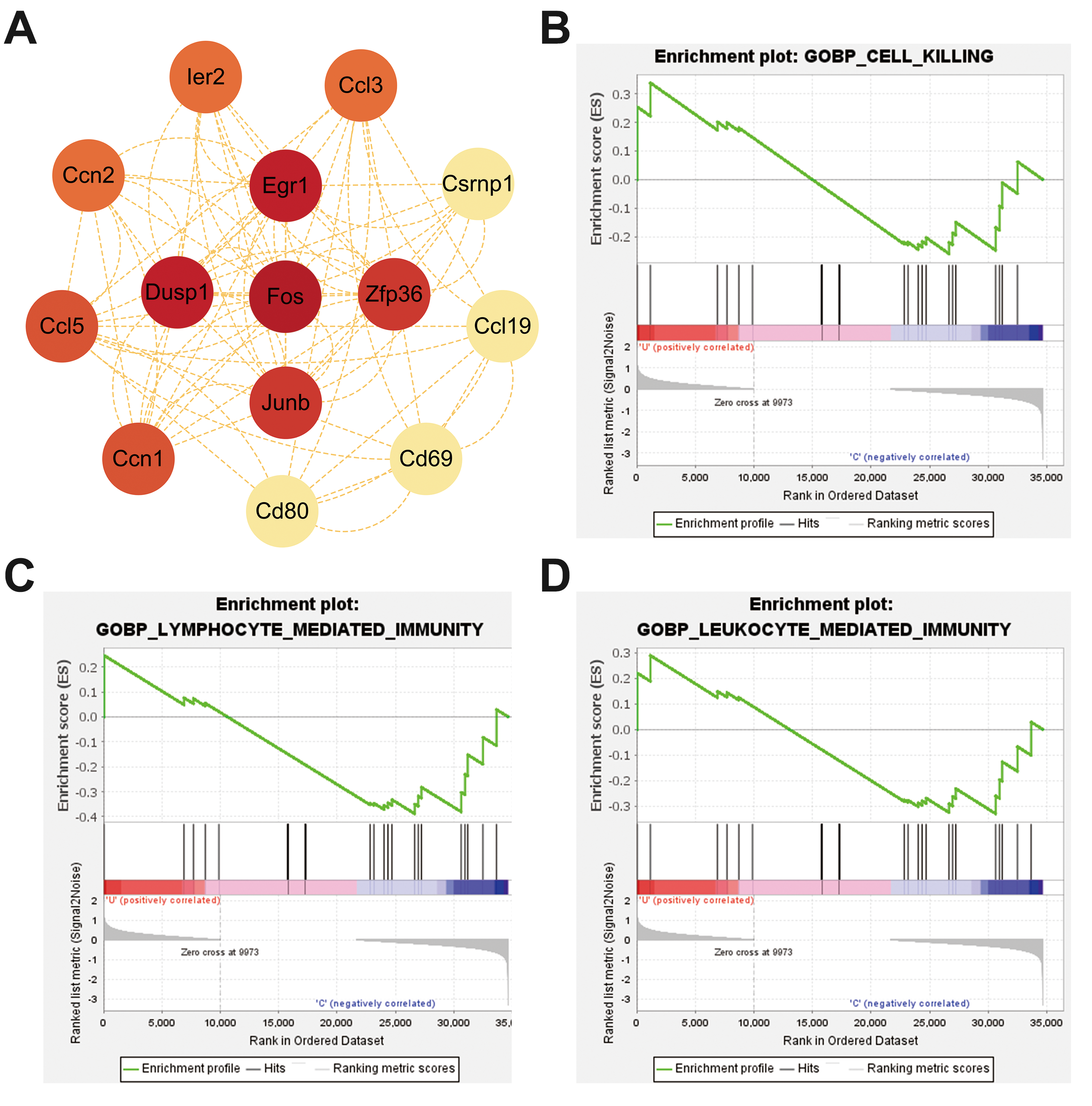

### Supplemental Figure 5

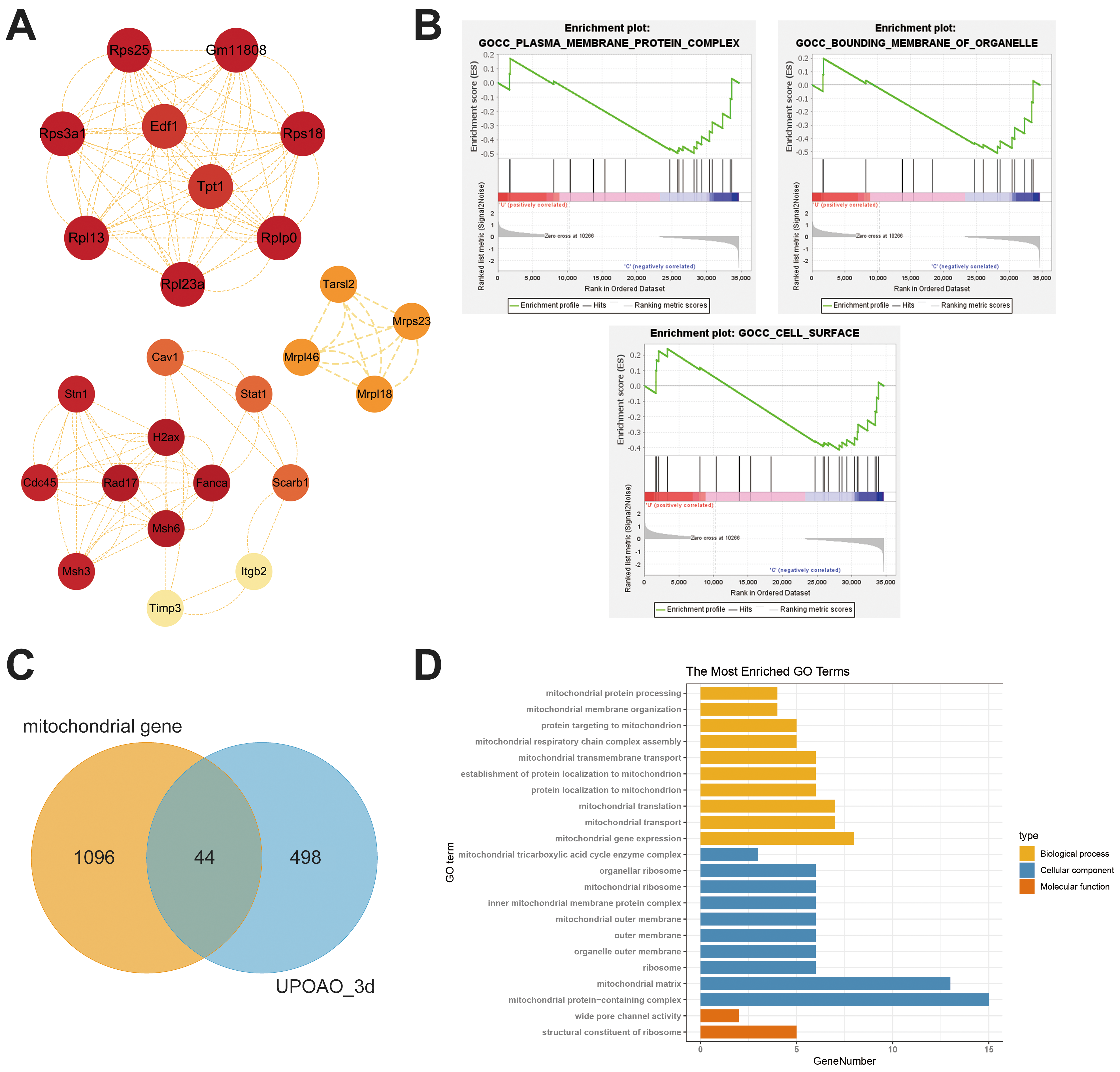

### Supplemental Figure 6

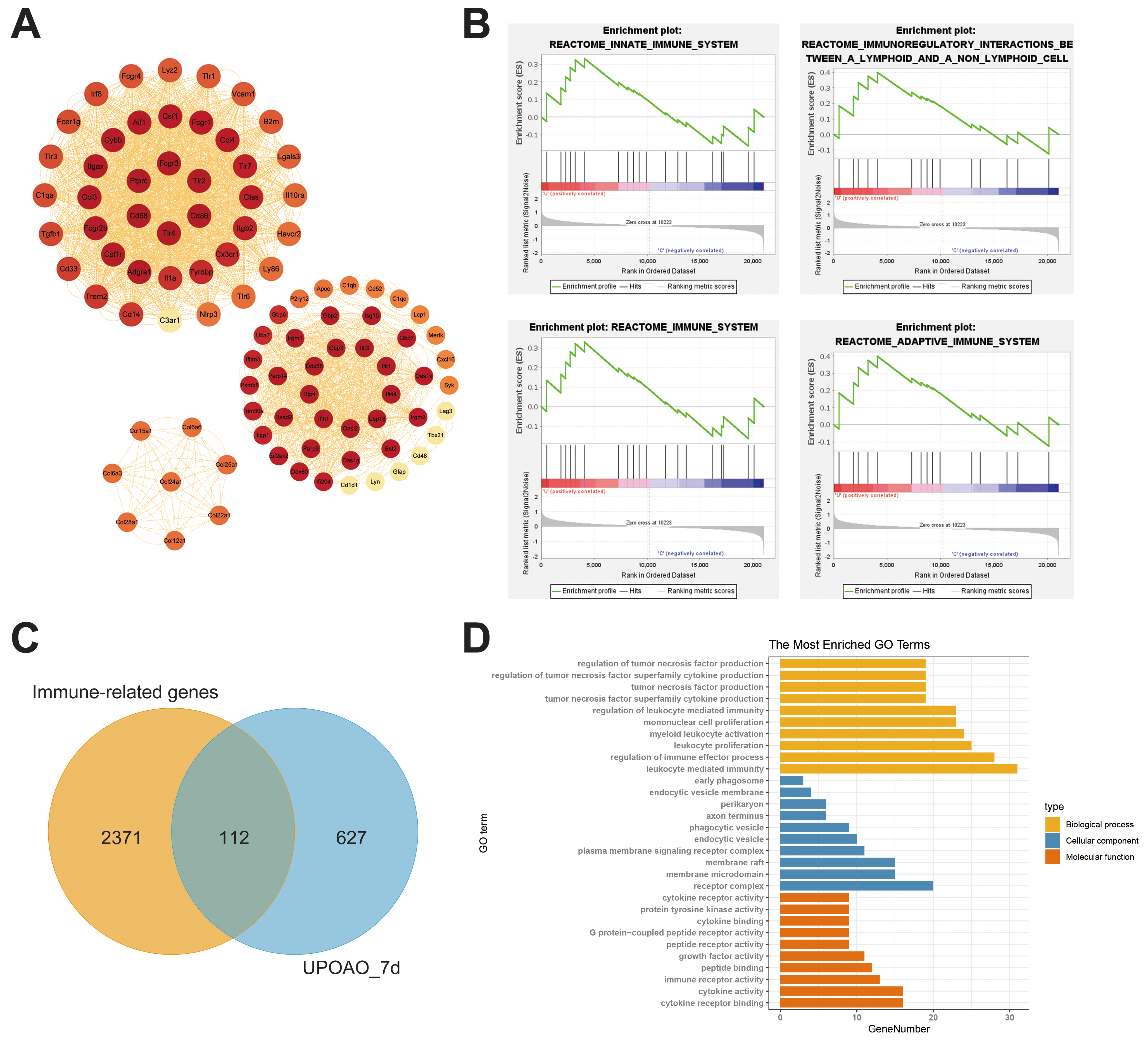

### Supplemental Figure 7

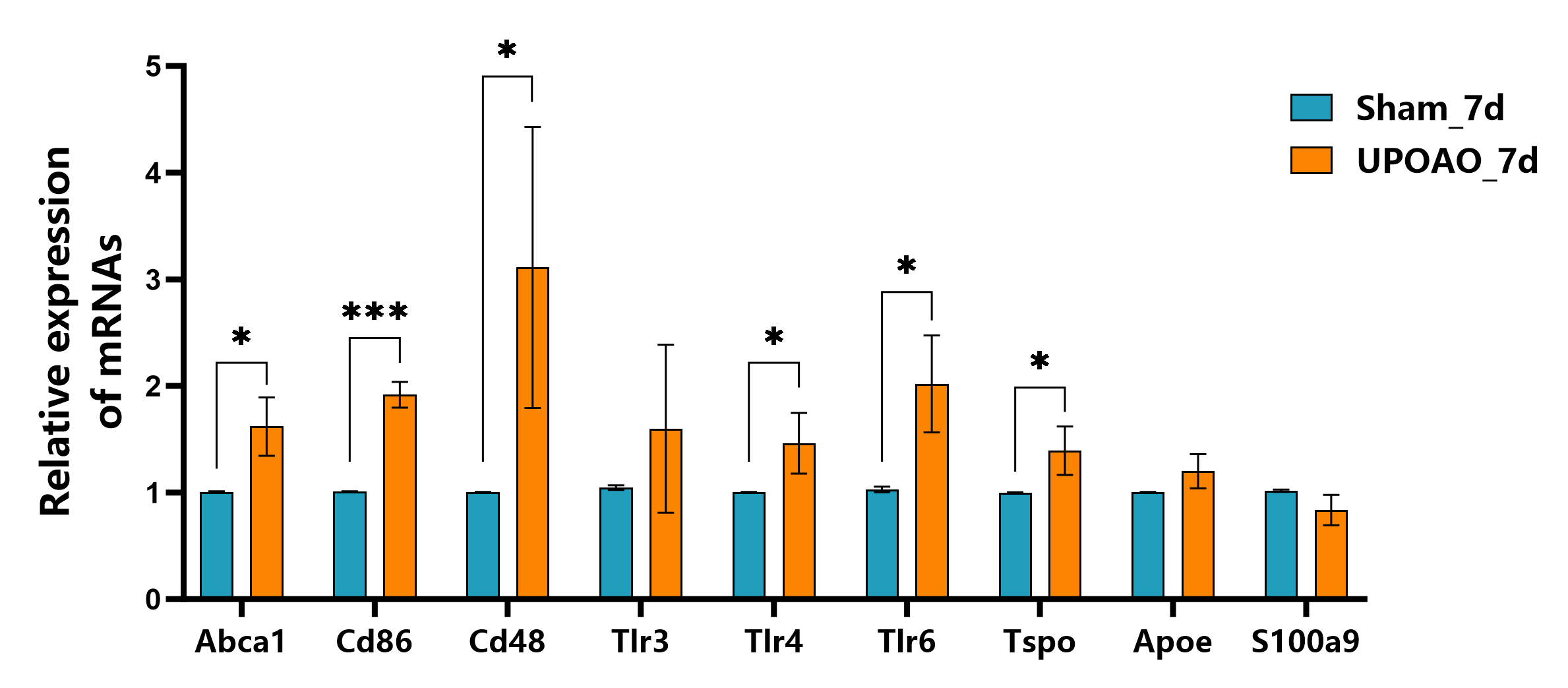

### Supplemental Figure 8

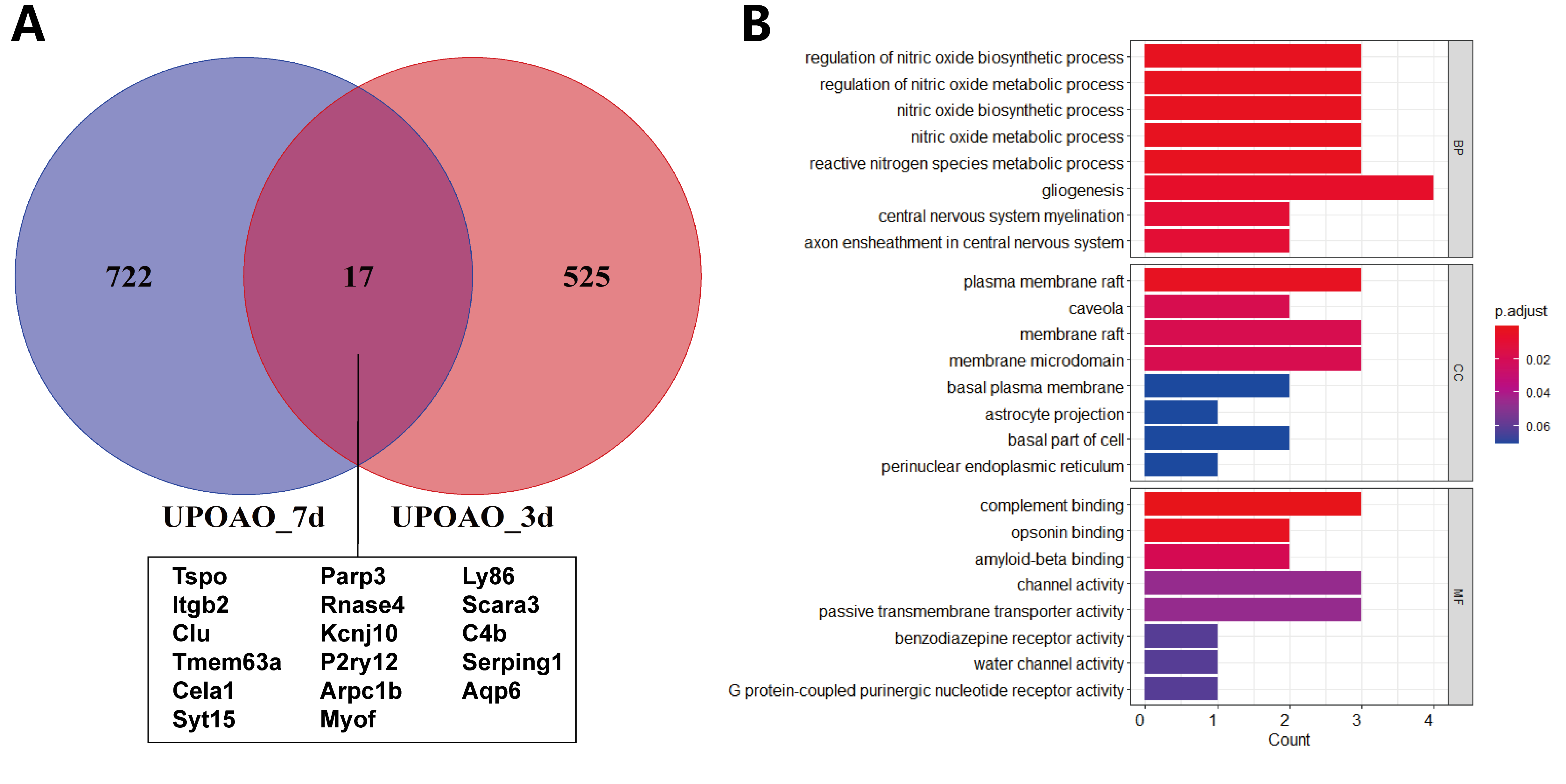
