## Supplemental Table 2 for "Silicone Wire Embolization-induced Acute Retinal Artery Ischemia and Reperfusion Model in Mouse: Gene Expression Provides Insight into Pathological Processes"

**Table. S2 Summary table:** Time course of all the morphological, functional, cellular, and transcriptome changes in the UPOAO model.

| Changes in each group | | | 30min-3d | 30min-7d | 60min-0d | 60min-3d | | | 60min-7d | | |
| --- | --- | --- | --- | --- | --- | --- | --- | --- | --- | --- | --- |
| RGCs counts | | | - | - | / | ↓ | | | ↓ | | |
| ERG | Amplitudes of a-waves | | - | - | / | - | | | - (0.001, 0.003, 0.01, 0.03, 0.1, 1.0 cd.s/m2)  ↓ (0.3, 3.0, 10.0 cd.s/m2) | | |
|  | Response time of a-waves | | - | - |  | - | | | - | | |
|  | Amplitudes of b-waves | | - | - |  | ↓ | | | ↓ | | |
|  | Response time of b-waves | | - | - |  | - | | | - | | |
|  | Amplitudes of OPs-waves | | / | | | ↓ | | | ↓ | | |
| OCT |  | | / | | | 1.5 PD | 3 PD | 4.5PD | 1.5 PD | 3 PD | 4.5PD |
|  | GCC thickness | |  |  |  | - | - | - | ↓ | ↓ | ↓ |
|  | INL+OPL thickness | |  |  |  | - | - | - | - | - | - |
|  | ONL+IS/OS+RPE thickness | |  |  |  | - | - | - | - | - | - |
|  | total retinal thickness | |  |  |  | - | - | ↓ | ↓ | ↓ | ↓ |
| HE | RNFL+GCL thickness | | / | | | ↑ | | | ↓ | | |
|  | IPL thickness | |  |  |  | ↓ | | | ↓ | | |
|  | INL thickness | |  |  |  | - | | | ↓ | | |
|  | OPL thickness | |  |  |  | - | | | - | | |
|  | ONL thickness | |  |  |  | - | | | - | | |
|  | IS/OS+RPE thickness | |  |  |  | - | | | - | | |
| Immunofluorescence density of retinal neural cells | Bipolar cells (PKCα) | | / | | | - | | | ↓ | | |
|  | Horizontal cells (Calbindin) | |  |  |  | ↓ | | | ↓ | | |
|  | Cholinergic amacrine cells (ChAT) | |  |  |  | ↓ | | | ↓ | | |
|  | Photoreceptor cells (Recoverin) | |  |  |  | - | | | - | | |
| Peripheral leukocyte infiltration (CD45) # | | | / | | | ↑ | | | ↑ | | |
| Microglia counts (Iba1) | | | / | | | ↑ | | | ↑ | | |
| RNA-seq | | DEGs number | / | | 419 (up: 215; down: 204) | 542 (up: 372; down: 170) | | | 739 (up: 429; down: 310) | | |
|  |  | Enrichment |  |  | Immune cells migration-related pathways | Oxidative stress-related pathways | | | Immune inflammation-related pathways | | |

-: no change;

↑: increase;

↓: decrease;

/: not involved in the article;

#: supplementary content: also increase after 60min-ischemia and 1d-reperfusion;
